# The mitotic CIP2A-TOPBP1 axis facilitates mitotic pathway choice between MiDAS and MMEJ

**DOI:** 10.1101/2024.11.12.621593

**Authors:** Peter R Martin, Jadwiga Nieminuszczy, Zuza Kozik, Nihal Jakub, Maxime Lecot, Julia Vorhauser, Karen A Lane, Alexandra Kanellou, Jörg Mansfeld, Laurence H Pearl, Antony W Oliver, Jessica A Downs, Jyoti Choudhary, Matthew Day, Wojciech Niedzwiedz

**Affiliations:** Division of Cancer Biology, The Institute of Cancer Research; London, SW3 6JB, UK; Faculty of Medicine of Rennes, University of Rennes, Rennes; F-35043, France; Division of Structural Biology, The Institute of Cancer Research; London, SW3 6JB, UK; Genome Damage and Stability Centre, University of Sussex; Brighton, BN1 9RQ, UK; Centre of Molecular Cell Biology, Queen Mary University of London; London, E1 4DQ, UK

## Abstract

Mitotic DNA double-strand breaks (DSBs) accumulate in response to replication stress or BRCA1/2 deficiency posing a significant threat to genome stability as repair by non-homologous end-joining (NHEJ) and homologous recombination (HR) is inactivated in mitosis. Mitotic cells instead rely on the mechanisms of microhomology mediated end-joining (MMEJ) and mitotic DNA synthesis (MiDAS). However, how these pathways are regulated in mitosis remains unknown. Here we reveal the CIP2A-TOPBP1 complex facilitates recruitment of SMX complex components to mitotic chromatin marked by CIP2A, through a CDK1-dependent interaction between TOPBP1 BRCT 1/2 and SLX4 phospho-threonine1260, that drives MiDAS. Furthermore, CIP2A promotes the recruitment of Polθ to facilitate mitotic MMEJ. This defines a mechanistic framework for mitotic DSB repair, where simultaneous disruption of MiDAS and MMEJ pathways underpins the synthetic lethality observed in BRCA1/2-deficient cells following CIP2A depletion. These findings provide critical insights into mitotic DNA repair and highlights therapeutic opportunities in HR deficient tumours.

## Introduction

DNA double-strand breaks (DSBs) in mitotic cells may persist from interphase due to replication stress or may occur directly in mitosis ^1^. If left unrepaired, these DSBs lead to gross chromosome breakage, genomic rearrangements, and polyploidy, all hallmarks of cancer ^2–5^. Established anti-cancer therapeutic approaches exploit this by induction of replication associated DNA damage, driving cells with unrepaired DNA into mitosis, promoting mitotic catastrophe and cell death^1, 6–9^. However, clinical resistance to these therapeutic modalities is a growing issue and underscores the gap in our mechanistic understanding of DNA repair outside of interphase.

Cells deficient in homologous recombination (HR), experience heightened replication stress, and are particularly prone to the accumulation of mitotic DNA double-strand break (DSBs). Recent studies have identified the role of DNA polymerase theta (Polθ) mediated microhomology-mediated end joining (MMEJ) and mitotic DNA synthesis (MiDAS) as critical repair pathways in mitosis, functioning redundantly to HR ^5, 10–12^. These pathways present a targetable vulnerability in tumours that exhibit heightened DNA replication stress, including those that are HR deficient ^1^. Despite the potential to target the mechanisms of DSB repair in mitosis, they remain inadequately defined, limiting our ability to fully exploit these mitotic mechanisms for anti-cancer therapy ^1, 6–8, 13–17^.

The SMX tri-nuclease complex is essential for genome stability and is activated in a phosphorylation-dependent manner specifically at the G2/M boundary ^18^. It plays an evolutionary conserved role in resolving replication and recombination intermediates and facilitating MiDAS to ensure proper chromosomal segregation ^11, 12, 18, 19^. SLX4 serves as a central hub coordinating the assembly and activity of SMX components at DNA damage sites ^18, 20^. Mutations in SLX4 (FANCP) cause Fanconi anemia, a disorder characterised by developmental abnormalities, anemia, and increased cancer risk ^21–23^. Its function is tightly regulated by kinases to prevent premature activation, as uncontrolled activity—such as Cyclin-dependent kinase 1 (CDK1) de-repression by Wee1-like protein kinase (WEE1) inhibition—promotes aberrant cleavage of replication forks, leading to chromosomal damage^20^. Together, this underscores the importance of the SMX complex and its regulation in the maintenance of genome stability.

TOPBP1, is an essential adaptor protein, with key roles in the maintenance of genome stability, largely attributed to its nine BRCT domains, which facilitate phosphorylation-dependent protein-protein interaction throughout the cell cycle, including in mitosis ^24–27^. Its functions are closely regulated by CDK1 and Polo-like kinase 1 (PLK1)-dependent phosphorylation events ^5, 28–31^. Notably, TOPBP1 interacts with DNA polymerase theta (Polθ), facilitating MMEJ-mediated double-strand break (DSB) repair during mitosis ^10^. TOPBP1 also plays a role in regulating the BTR (BLM-TOP3a-RMI1/2) dissolvase complex, supporting chromosomal segregation through phosphorylation events driven by CDK1 and PLK1^30^. Studies have also shown that the interaction between yeast orthologs of TOPBP1 (Dbp11) and SLX4 (Slx4) is mediated by CDK1-driven phosphorylation^28^, a mechanism that is conserved in human cells, where the interaction is facilitated through SLX4 phosphorylation at Thr1260 ^28, 29^. However, the role of the TOPBP1-SLX4 interaction in mitotic repair especially in context of HR deficiency remains undefined.

Recently a mitotic specific DNA repair complex consisting of MDC1, TOPBP1 and CIP2A has been identified which functions to tether broken chromosomes to prevent fragmentation and mitotic catastrophe ^32, 33^. This mechanism is independent of Polθ, POLD3, or the MRN complex components, although their loss increases mitotic DNA damage ^32^. CIP2A and TOPBP1, interact directly and exhibit mutual dependence for chromatin recruitment during mitosis ^13^. Recent studies have underscored CIP2A and its interaction with TOPBP1 as a highly penetrant mitotic specific synthetic lethal (SL) target in BRCA1/2 deficient cells^1, 13^. The mechanistic underpinning of this synthetic lethal interaction is unknown; although, it has been suggested that CIP2A’s role in the mitotic DNA tethering complex may be a contributing factor ^8, 32, 33^. However, this explanation is challenged by the observation that MDC1 loss, another key DNA tethering factor does not induce the same synthetic lethality in BRCA1/2 deficiency^34, 35^. An alternative, hypothesis is that CIP2A is a key regulator of yet undefined mitotic DNA repair. However, although the interaction between TOPBP1 and Polθ is essential for MMEJ-dependent DSB repair in mitosis^5^, CIP2A appears to be dispensable for MMEJ-driven telomere fusions, even though it is critical in regulating TOPBP1 recruitment to chromatin ^1, 13, 36, 37^. This suggests that other targetable factors/mechanisms may drive the synthetic lethality between CIP2A-TOPBP1 and BRCA1/2 deficiency. Given the limited understanding of mitotic DSB repair mechanisms and the ongoing clinical trials targeting Polθ, defining the precise role of the CIP2A-TOPBP1 axis in DNA repair is critical for improving the treatment of cancers.

To address this, we conducted comprehensive unbiased Co-IP/MS analysis, that mapped the TOPBP1 interactome across different cell cycle phases. Importantly, we identify key interactions in mitosis with SMX complex components (SLX4, ERCC1, XPF, EME1, MUS81), CIP2A, MDC1, and PLK1. We determine that TOPBP1 is required for the recruitment of MUS81 in BRCA2 deficiency. We identify CDK1 dependent phosphorylation of SLX4 at Thr1260 as critical for its interaction with BRCT1 and 2 of TOPBP1. This interaction functions to recruit SLX4, MUS81 and ERCC1 to mitotic chromatin marked by the CIP2A-TOPBP1 complex in response to replication stress. Cells with a Thr1260Ala mutation show defective MiDAS, leading to increased genome instability, indicated by elevated micronuclei levels.

Critically, our study uncovers a genetic interaction network that defines the CIP2A-TOPBP1 complex as a master regulator that facilitates mitotic DNA repair pathway choice between MiDAS and MMEJ. The CIP2A-TOPBP1 complex achieves this by recruiting not only components of the SMX complex (SLX4, MUS81 and ERCC1) but also Polθ to mitotic chromatin. Concurrently, we demonstrate, that CIP2A loss impairs both break induced replication (BIR)-lke /MiDAS and MMEJ. Pharmacological inhibition of Polθ, combined with loss of the TOPBP1-SLX4 interaction, exacerbates genome instability and reduces cellular proliferation under replication stress. Notably, SLX4, Polθ, and CIP2A are essential for cellular proliferation in BRCA1/2-deficient cells and expression of a minimal SLX4 fragment containing Thr1260 is also sufficient to impair proliferation in BRCA1/2 deficient cells. This study elucidates the central role of the CIP2A-TOPBP1 complex in the control of mitotic DNA repair, providing a mechanistic framework that underpins its synthetic lethal interaction with BRCA1/2 mutations. Consequently, these findings further emphasise the CIP2A-TOPBP1 axis as a promising therapeutic target in BRCA1/2-deficient cancers and those with elevated DNA replication stress, while highlighting new actionable avenues for anti-cancer therapy development.

## Results

### Cell cycle-specific proteomics reveals TOPBP1 interactors in mitosis

TOPBP1 is frequently overexpressed in multiple cancers, including breast cancer and its overexpression in this setting is associated with an aggressive phenotype^38–40^. Moreover, single-nucleotide polymorphisms (SNPs) in *TopBP1* resulting in higher protein expression have been associated with an increased risk in breast and endometrial cancers^41, 42^. These tumours are heavily burdened with BRCA1/2 mutations, resulting in significant homologous recombination deficiency (HRD) ^43–45^. With growing interest in exploiting mitotic double-strand break (DSB) repair as a therapeutic vulnerability in HR deficient tumours, we hypothesised that the function of TOPBP1 in mediating protein-protein interactions throughout the cell cycle may be especially critical during mitosis ^1, 5, 10^. To explore this, we conducted an unbiased, cell-cycle stage-specific, co-immunoprecipitation coupled with LC-MS analysis to identify proteins uniquely enriched during S-phase and mitosis (Fig.1A, B, C and Fig. S1A). This approach allowed us to map the mitotic TOPBP1 interactome, revealing proteins preferentially enriched in M-phase compared to asynchronous or S-phase cells (Fig. 1B and C), including previously reported factors PLK1, MDC1, and CIP2A ^1, 13, 37, 46^. Notably, we identified elevated interaction between TOPBP1 and members of the SMX complex in mitosis. These findings reveal dynamic changes in TOPBP1 complexes throughout the cell-cycle and provide new insights into its role in mitosis.

**Fig. 1:**
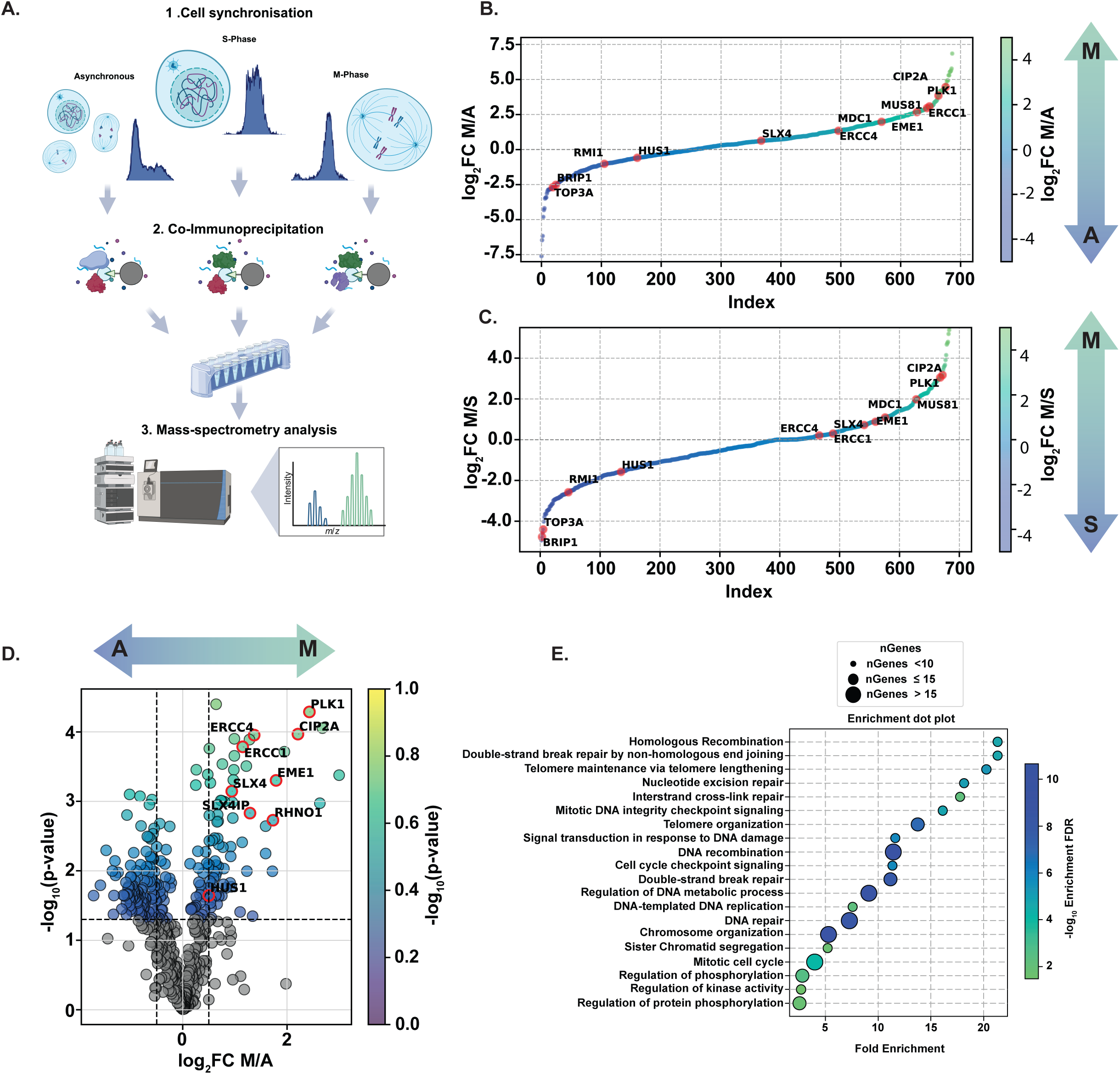
Enrichment of TOPBP1 interactions with SMX complex components during mitosis. (A) Schematic representation of endogenous TOPBP1 Co-immunoprecipitation experiments across different cell cycle phases from independent HEK293TN cells (created with BioRender.com). (B). Dot plot illustrating mean log_2_ fold change (FC) of proteins detected in TOPBP1 Co-IP samples from M phase versus asynchronous (A) cells, by-MS label-free mass-spectrometry. (C) similar analysis as in B, comparing S phase with M phase. Samples analysed in A, B and C were from three independent experiments. (D) Dot plot representing mean of TMT quantitative mass spectrometry analysis showing statistically significant enrichment of TOPBP1 interactions in asynchronous versus M phase cells, samples were from three independent experiments. Statistical significance was determined by two tailed t-test. (E) Gene Ontology (GO)-term enrichment analysis of the mitotic TOPBP1 interactome −log10 enrichment false discovery rate (FDR).

To further validate and explore the functional implications of the TOPBP1 interactome, we performed an additional quantitative Tandem Mass Tag (TMT) mass-spectrometry analysis by isobaric labelling of TOPBP1 Co-IPs from asynchronous and M phase synchronised cells (Fig. 1D). This analysis highlighted a subset of proteins significantly enriched in mitosis, including key DNA repair proteins, such as PLK1, CIP2A, RHNO1, EME1, SLX4, ERCC4 and SLX4IP, supporting the notion that TOPBP1 plays a crucial role in orchestration of a network of DNA repair complexes specifically during mitosis (Fig.1D).

To gain a broader understanding of the biological processes associated with the TOPBP1 interactome, we conducted a gene ontology (GO) enrichment analysis on the identified proteins in our TMT analysis (Fig. 1E). The analysis revealed a significant enrichment for processes related to DNA repair, mitotic cell cycle, chromosome organization, and protein phosphorylation. Notably, pathways such as double-strand break repair, homologous recombination, and non-homologous end joining were among the top enriched categories, supporting a role of TOPBP1 in promoting various aspects of mitotic DNA repair and maintenance of genome stability during cell division (Fig. 1E). Subsequently, to gain phenotypic insight into the role of TOPBP1 in supporting genome stability in mitosis we employed a TOPBP1-degron system ^47^. HCT116 cells expressing osTIR1 that were CRISPR edited at the endogenous locus to express TOPBP1-mAID-Clover were exposed to IAA to elicit acute degradation of TOPBP1. Consistent with the crucial role of TOPBP1 in mitosis, its acute temporal depletion in G2/M, followed by release into mitosis led to a marked increase in anaphase abnormalities, canonical markers of genome instability, including DAPI bridges, DNA laggards, and PICH-marked ultra-fine anaphase bridges (UFBs) (Fig. S1B, C and D). Taken together we hypothesise that TOPBP1 may play an essential role in mitotic genome stability through identified mitotic protein-protein interactions.

Notably, in our mass-spectrometry based interactome analysis, we observed significant enrichment of the majority of the components of the SMX complex (ERCC4/XPF, ERCC1, EME1, MUS81 and SLX4 (Fig1. B, C and D)) indicating a possible functional role of a TOPBP1-SMX complex interaction that may play a fundamental role in the maintenance of genome stability. Therefore, we sought to validate this interaction by eGFP-TOPBP1 co-immunoprecipitation from human cell extracts and found mitotic enrichment of endogenous components of the SMX complex (SLX4, MUS81 and ERCC1 (Fig. S1E; lysates were treated with benzonase to prevent DNA bridging)). Thus, we conclude that TOPBP1 and its interaction with components of the SMX complex is elevated in mitosis.

### TOPBP1 co-localises with SMX components and is required for MUS81 recruitment in mitosis

BRCA2 loss is associated with increased replicative stress and the accumulation of unresolved replication and recombination intermediates in mitosis^1^. To determine whether BRCA2 deficient cells demonstrate reliance on TOPBP1 associated functions we analysed TOBPP1 recruitment in mitotic cells. We indeed observed an increase in the incidence of TOPBP1 foci in prometaphase DLD1 BRCA2^−/−^ cells, indicating elevated requirement for mitotic TOPBP1 function in these cells (Fig. S2A). To gain insight into whether BRCA2 deficient cells demonstrate dependency on the association of TOPBP1 and SMX complex components in mitosis we analysed MUS81 and ERCC1 colocalisation with TOPBP1 in prometaphase DLD1 BRCA2^−/−^ cells (Fig. 2A and B). As such, TOPBP1 is recruited to chromatin and frequently co-localises with MUS81 and ERCC1 in prometaphase cells (Fig. 2A and B). Notably, we observed a significant elevation of both MUS81-TOPBP1 and ERCC1-TOPBP1 colocalisation events in BRCA2 deficient cells when compared to their WT control (Fig. 2A and B). Similarly, treatment of non-cancerous RPE1 cells with aphidicolin, to induce replicative stress, resulted in elevated mitotic colocalisation of MUS81 and ERCC1 with TOPBP1 in prometaphase (Fig. S2B and C). Indicating, TOPBP1 and components of the SMX complex associate in mitosis in response to replication stress or BRCA2 deficiency consistent with their defined roles in repair of under-replicated DNA or persistent recombination intermediates.

**Fig. 2:**
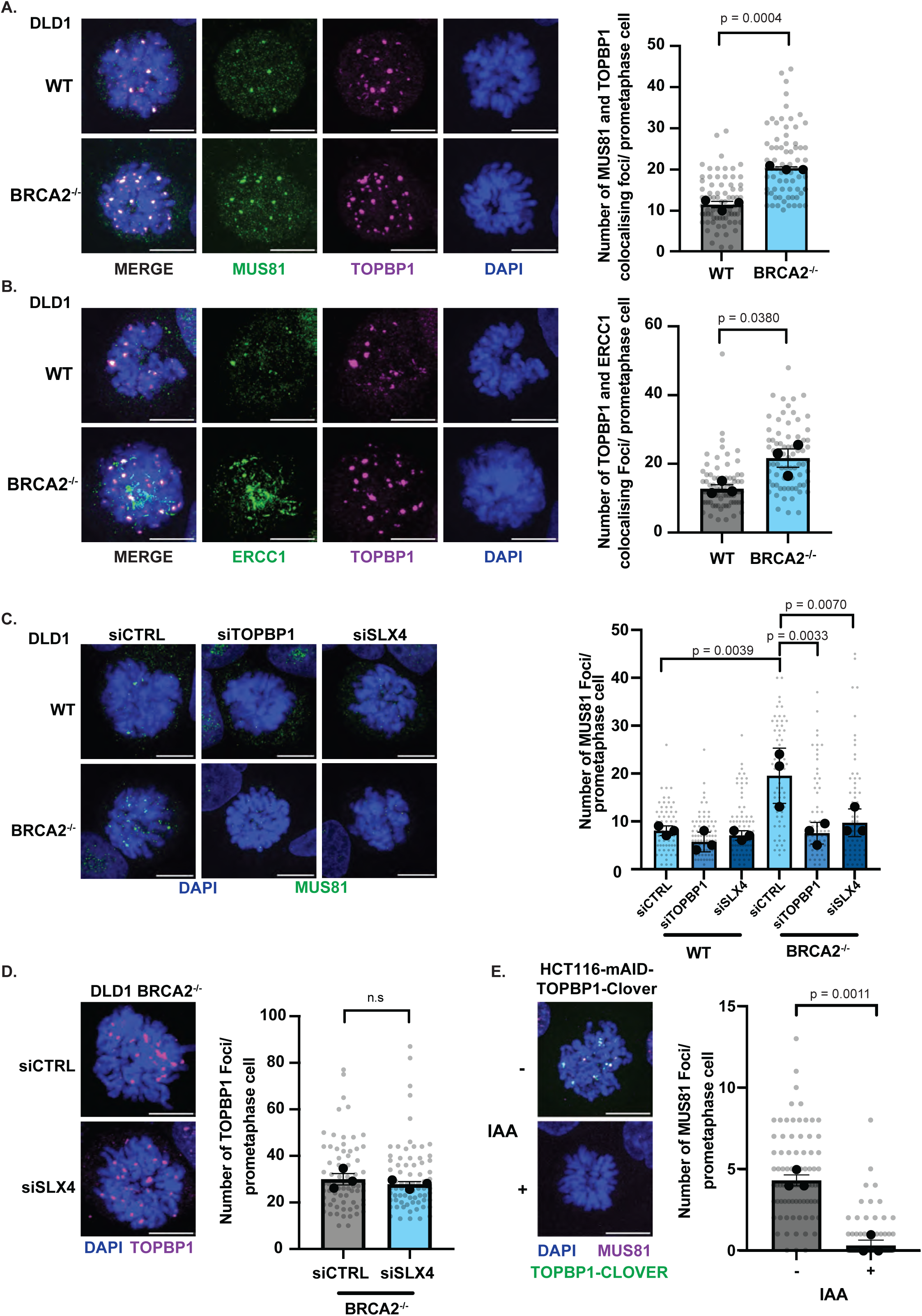
Elevation of mitotic TOPBP1 dependent recruitment of SMX complex components in BRCA2 deficiency. (A) Representative images and a dot plot showing the number of MUS81 and TOPBP1 colocalising foci in DLD1 and DLD1 BRCA2^−/−^ prometaphase cells (WT: n=74 and BRCA2^−/−^: n=73, from three independent experiments, statistical significance was). (B) Representative images and a dot plot showing the number of ERCC1 and TOPBP1 colocalising foci in DLD1 and DLD1 BRCA2^−/−^ cells (WT: n=73, BRCA2^−/−^: n=74, from three independent experiments). (C) Representative images and a dot plot showing the number of MUS81 localisation in DLD1 and DLD1 BRCA2^−/−^ prometaphase cells, treated with siCTRL, siTOPBP1 or siSLX4 (DLD1 (siCTRL: n=75, siTOPBP1: n=82, siSLX4: n=86) and DLD1 BRCA2^−/−^ (siCTRL: n=65, siTOPBP1: n=69, siSLX4: n=64) from three independent experiments). (D) Representative images and a dot plot showing TOPBP1 foci in DLD1 BRCA2^−/−^ prometaphase cells (siCTRL: n=72 and siSLX4: n=73, from three independent experiments). (E) Representative images and a dot plot showing number of MUS81 foci in prometaphase HCT116-TOPBP1-mAID-Clover cells with and without 2 hour IAA (500µM) dependent degradation of TOPBP1 in the presence of 40 ng/ml nocodazole (“-“: n=81 and “+”: n=84, from three independent experiments. Statistical significance in A, B, D and E was determined by two-tailed unpaired t-test. Statistical significance in C was determined by the two-way ANOVA with Tukey’s post-hoc test. In A-E individual measurements are represented by grey dots whereas black dots indicate medians of each experiment, and the mean is presented as bars with error bars showing S.E.M. Scale bars equivalent to 10 µm.

Given the observed elevation in MUS81-TOPBP1 colocalisation in BRCA2^−/−^ cells, we sought to ascertain the requirement of TOPBP1 and SLX4 in mediating MUS81 localisation in wild type and BRCA2^−/−^ DLD1 prometaphase cells (Fig. 2C). Consistently, TOPBP1 depletion by siRNA resulted in a marked reduction in MUS81 foci formation in both WT and BRCA2^−/−^ cells compared to siRNA control treated cells (Fig. 2C). Similarly, SLX4 depletion, resulted in defects in its localisation akin to those observed with TOPBP1 depletion (Fig. 2D, S2E and S2F). However, SLX4 depletion in BRCA2^−/−^ cells did not impair TOPBP1 localisation to prometaphase chromatin, suggesting SLX4 and MUS81 act downstream of TOPBP1 (Fig 2D). Together this data indicate that the recruitment of the SMX components to chromatin in mitosis is elevated in BRCA2 deficiency and in response to replication stress and are mediated by both TOPBP1 and SLX4.

To determine whether TOPBP1 is required specifically in mitosis for localisation of MUS81, we employed the HCT116-TOPBP1-mAID-Clover degron system to facilitate acute depletion of TOPBP1 by addition of IAA for 2 hours, followed by analysis of prometaphase cells ^47^ (Fig. S1D and S2D). In the presence of TOPBP1 we observed robust formation of MUS81 foci in prometaphase cells (Fig. 2E). In contrast, cells acutely depleted of TOPBP1 demonstrated a near total abolishment of MUS81 foci formation in prometaphase (Fig. 2E), demonstrating TOPBP1 is required in mitosis for MUS81 recruitment.

### The N-terminal module of TOPBP1, containing BRCT 0-1-2, mediates interaction with the SMX complex in a phospho-dependent manner

TOPBP1 consists of nine BRCT domains, which facilitate phosphorylation-dependent protein-protein interactions in a number of distinct complexes ^24, 27^ (Fig. 3A). To gain insight into the mitotic function of TOPBP1, we sought to map binding site(s) mediating its interaction with components of the SMX complex. To achieve this, we expressed C-terminal truncation mutants of TOPBP1 in mitotic synchronised HEK293TN cells and performed co-immunoprecipitations (Fig. 3B and S3A). Strikingly, all TOPBP1 truncation mutants tested, including BRCT0-2, were sufficient to bind SLX4, MUS81 and ERCC1 of the SMX complex (Fig. 3B). However, interaction between TOPBP1 and the BTR complex components BLM and TOP3a were significantly reduced in cells expressing the BRCT 0-2 and 0-3 mutants, consistent with their previously characterised binding site in BRCT 5 of TOPBP1 (Fig. 3B)^31^. Importantly, N-terminal truncation of TOPBP1, that removed BRCT 0-2 but maintained all other domains of TOPBP1, was sufficient to disrupt interaction between TOPBP1 and components of the SMX complex in mitotic synchronised cells (Fig. 3C). Therefore, BRCT 0-2 of TOPBP1 is both necessary and sufficient to mediate its interaction with components of the SMX.

**Fig. 3:**
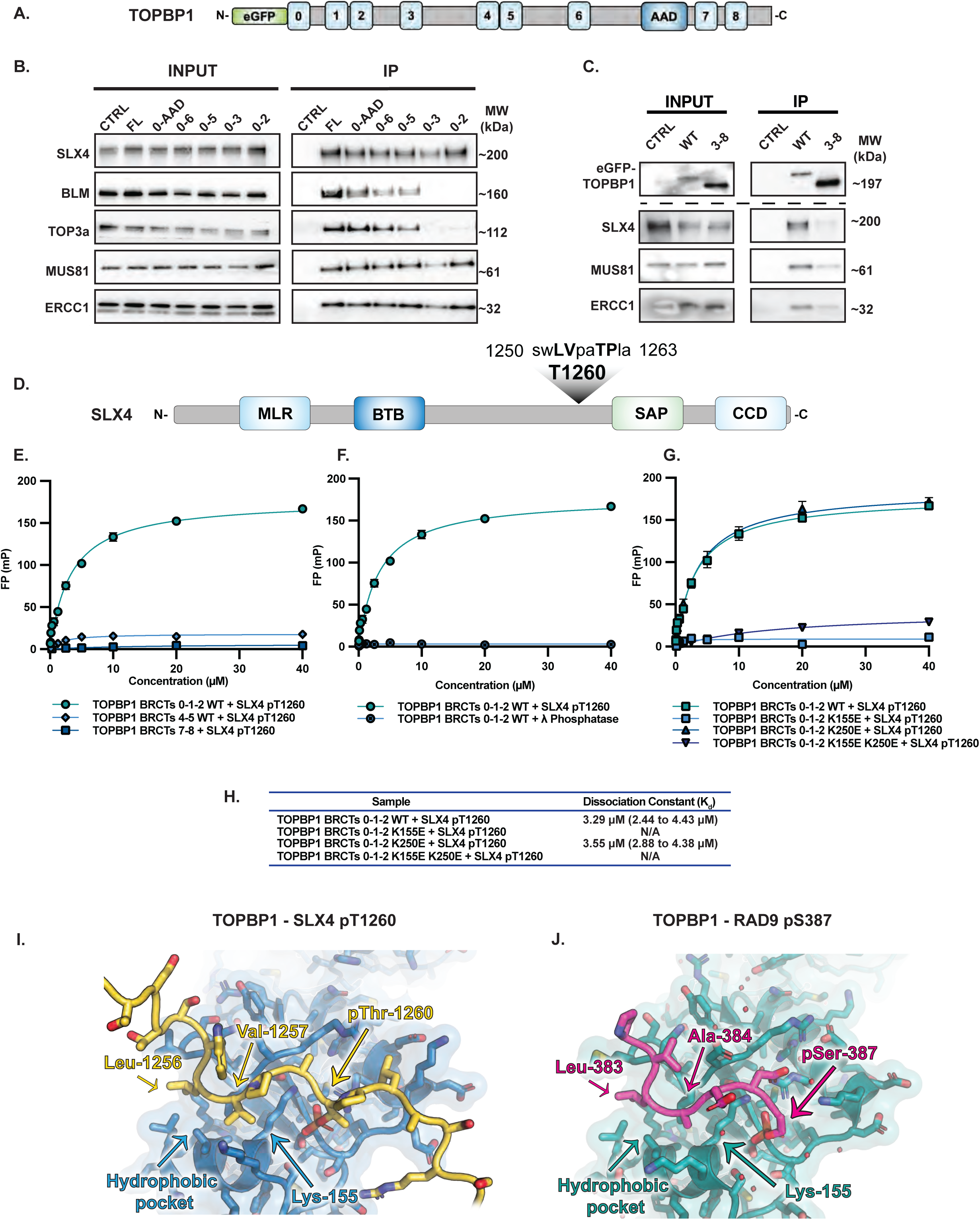
SLX4 interacts with BRCT1/2 of TOPBP1 via pT1260 in mitotic cells. (A) eGFP-TOPBP1 domain architecture schematic (B) Western blot analysis of eGFP-TOPBP1 interacting SMX complex proteins in eGFP-TOPBP1 truncation mutant Co-IP from HEK293TN cells transiently transfected with WT or C-terminally truncated eGFP-TOPBP1 expression constructs followed by 16 hours 100 ng/ml nocodazole synchronisation from Fig S3A. Mock Co-IP from non-transfected HEK293TN cells were used as negative control (CTRL). (C) Western blot analysis of SMX component interactions with TOPBP1 in eGFP-TOPBP1 Co-IPs from HEK293TN cells transiently transfected with WT or N-terminally truncated Δ314 (3-8) eGFP-TOPBP1 expression constructs followed by 16 hours 100 ng/ml nocodazole synchronisation. Mock Co-IP from non-transfected HEK293TN cells were used as negative control (CTRL). (D) Domain architecture of SLX4 and relative position of the identified T1260, TOPBP1 BRCT 1 recognition motif. (E) Fluorescence polarisation analysis of recombinant TOPBP1 BRCT0/1/2, BRCT 4/5 and BRCT 7/8 domain containing fragments in the presence of a fluorescein tagged SLX4 pT1260-containing peptide. For all IP experiments 1% of input was used for analysis by western blot of input lysate. (F) Fluorescence polarisation analysis of recombinant TOPBP1 BRCT0/1/2 fragment in the presence of fluorescein tagged SLX4 pT1260-containing peptide with or without lambda (λ) phosphatase treatment. (G) Fluorescence polarisation analysis of recombinant TOPBP1 WT or conserved lysine to glutamic acid mutations in BRCT1 (K155E) or 2 (K250E) or in BRCT 1 +2 (K155E +K250E) fragment in the presence of fluorescein tagged SLX4 pT1260 containing peptide. (H) Table showing the dissociation constant of TOPBP1 BRCT 0-1-2 WT or K250E with SLX4 pT1260. (I) TOPBP1 BRCT 0-1-2 and SLX4 pT260 interaction modelled in AlphaFold 3. (J) TOPBP1 BRCT 0-1-2 and RAD9 pS387 crystal structure (PDB: 6HM5).

Based on these observations, we hypothesised that the TOPBP1-SLX4 interaction may be direct and a canonical phosphorylation dependent – BRCT interaction ^25, 26^. Bioinformatic analysis of the SLX4 sequence, scanning for defined BRCT1 binding consensus motifs ‘[VIL]-[AVIL]-x-x-[ST]-P’ ^26^ identified two CDK consensus motifs centered on the highly conserved residues Thr1260 and Thr1476 (Fig 3D and S3B). Notably, phosphorylation of SLX4 Thr1260 and Thr1476 have been previously identified in human cells ^48^.

Using purified TOPBP1 protein fragments ^26^, we conducted *in vitro* fluorescence polarisation analysis in the presence of SLX4 peptides containing phospho-Thr1260 (pThr1260) and phospho-Thr1476 (pThr1476). We observed that the pThr1260-phosphorylated SLX4 peptide, interacted with a GST-tagged TOPBP1 fragment containing BRCTs 0-1-2 while the Thr1476 containing phosphorylated peptide did not bind (Fig.3E and Fig. S3B). Furthermore, no interaction was detected between TOPBP1 fragments containing BRCTs 4-5 or 7-8 for either the SLX4-pThr1260 and -pThr1476 peptides (Fig. 3E and Fig. S3B). Treatment of the pThr1260 peptide with lambda phosphatase prevented the interaction with BRCT0-1-2 protein (Fig. 3F), demonstrating the interaction is specific and phosphorylation-dependent.

Tandem BRCT domains contain conserved lysine residues that interact with the phosphate group of a modified protein ligand, and mutating this lysine drastically reduces ligand-binding affinity ^24, 26, 27, 37^. In TOPBP1, Lys155 in BRCT1 and Lys250 in BRCT2 are highly conserved. Introducing K155E, or K155E + K250E, mutations within BRCTs 1 and 2, disrupted the interaction between the TOPBP1 BRCT 0-1-2 containing fragment and the phosphorylated SLX4 pThr1260 peptide *in vitro* while the K250E mutant was able to bind with comparable K_d_ to WT (Fig. 3G and H). This demonstrated a preference of the pThr1260 peptide for BRCT1 of TOPBP1. Modelling of the TOPBP1 BRCT 0-1-2 SLX4 pThr1260 interaction in Alphafold 3 supported the structural basis of the interaction (Fig. 3I). Additionally, when comparing the modelled interaction to the previously characterised crystal structure of the TOPBP1 BRCT 0-1-2 and RAD9 pS387 interaction^26^, we observed strong similarities in BRCT1 dependent phospho-ligand recognition (Fig. 3I and J). As such, the phosphate group of SLX4 pThr1260 is predicted to be recognised by the conserved triplet residues Thr114, Arg121 and Lys155 in the BRCT1 domain of TOPBP1, as seen experimentally for the RAD9 pSer387 interaction (Fig. 3I and J). Furthermore, several of the interactions predicted N-terminal of the phosphate group residues are conserved with the hydrophobic side chains of SLX4 Leu1256 and Val1257, taking the place of RAD9 residues Leu383 and Ala384, seen sitting in the hydrophobic pocket on the surface that determines the specificity of BRCT1 (Fig. 3I and J) ^26^. Together, this provides a putative structural basis of the TOPBP1 BRCT 0-1-2 – SLX4 pThr1260 interaction.

To further validate the human TOPBP1 BRCT1 interaction with SLX4 pThr1260, we expressed eGFP-tagged full-length TOPBP1 containing lysine-to-glutamic acid mutations in BRCT domains 1, 2, and combined 1 and 2. We expressed these proteins in mitotic synchronised cells and carried out co-immunoprecipitation analysis (Fig. 4A). We validated this approach by probing for MDC1, a known TOPBP1-binding protein that interacts with BRCTs 1 and 2 ^24^. As expected, mutation of BRCT1 or BRCT2 specifically prevented MDC1 binding (Fig. 4A) ^13^.

**Fig. 4:**
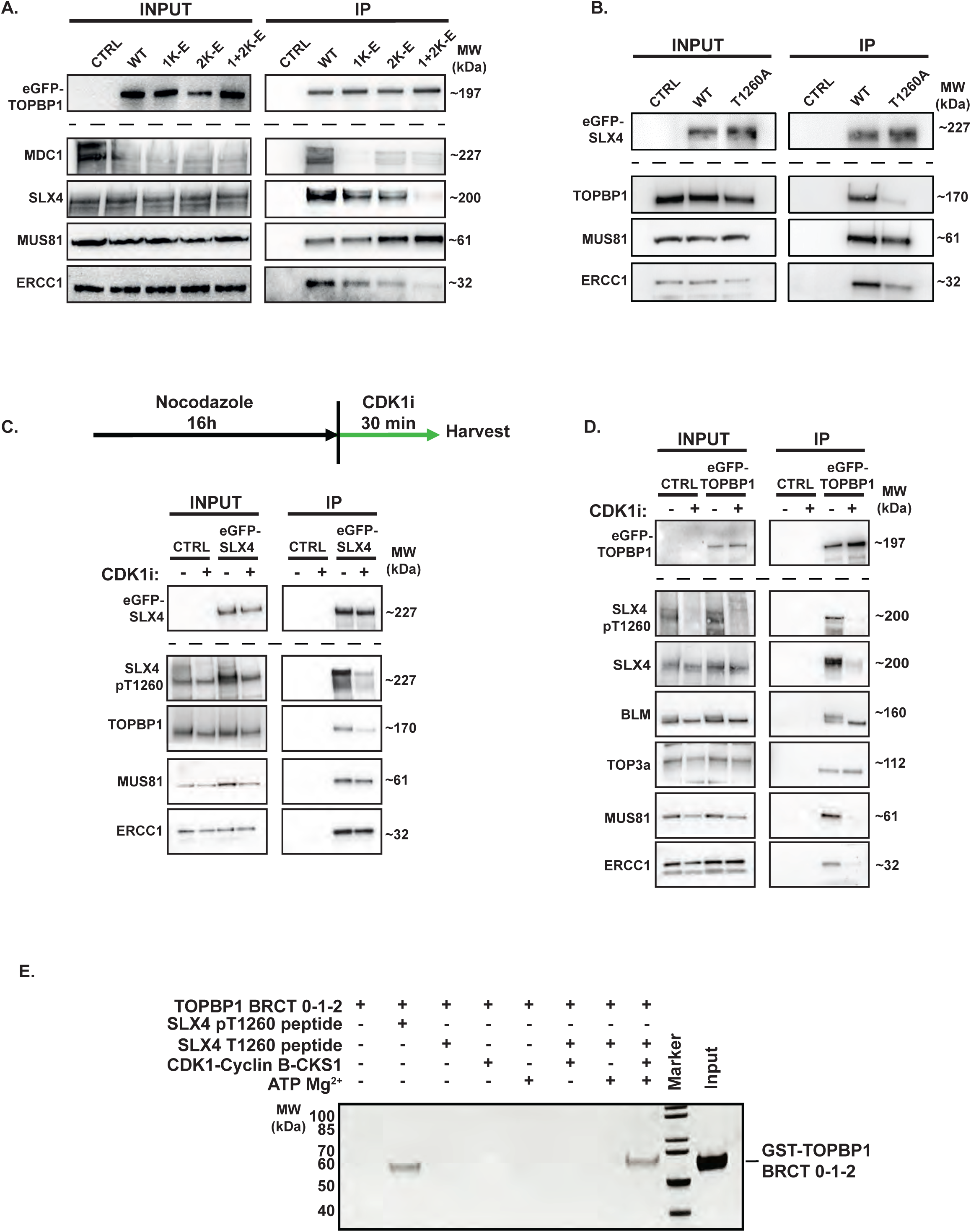
Mitotic interaction of the SMX complex components and TOPBP1 is driven by CDK1 activity. (A) Western blot analysis of MDC1 and SMX complex component interaction with eGFP-TOPBP1 in Co-IPs from HEK293TN cells treated with 100 ng/ml of nocodazole for 16h after transient transfection of eGFP-TOPBP1 WT, BRCT 1 (K155E), 2 (K250E) or 1+2 (K155E +K250E) expression constructs. Mock Co-IP from non-transfected HEK293TN cells were used as negative control (CTRL). (B) Western blot analysis of TOPBP1 and SMX component interaction with eGFP-SLX4 in Co-IPs from HEK293TN cells treated with 100 ng/ml of nocodazole for 16h after transient transfection of eGFP-SLX4 WT and T1260A expression constructs. Mock Co-IP from non-transfected HEK293TN cells were used as negative control (CTRL). (C) Western blot analysis of CDK1 dependent phosphorylation and interaction between SMX components and TOPBP1 in eGFP-SLX4 Co-IPs. HEK293TN cells were transfected with an eGFP-SLX4 expression construct and treated with or without 9 µM RO-3306 (CDK1i) 30 minutes after 16h 100 ng/ml nocodazole treatment. Mock Co-IP from non-transfected HEK293TN cells act as negative control (CTRL). (D) As in C, but reciprocal eGFP-TOPBP1 Co-IP analysis. For all IP experiments 1% of input was used for analysis by western blot of input lysate. (E) SDS-PAGE of *in vitro* reconstitution of CDK1-Cyclin B-CKS1 driven phosphorylation of a SLX4 T1260 containing biotinylated peptide and its phosphorylation-dependent interaction with a recombinant TOPBP1 BRCT 0-1-2 containing fragment.

We next assessed the ability of these TOPBP1 mutants to interact with SLX4, MUS81 and ERCC1. Consistent with our fluorescence polarisation data, we found that the K155E/K250E double mutant, which disrupts both BRCT1 and BRCT2, impaired the ability of TOPBP1 to bind SLX4 and ERCC1 (Fig. 4A). This suggests that while BRCT1 is critical for the interaction, BRCT2 likely provides additional stabilising interactions or facilitates binding indirectly of TOPBP1 or SLX4. Strikingly, MUS81 binding was unaffected by the disruption of BRCT1 and BRCT2 (Fig. 4A), highlighting a TOPBP1 BRCT1/2-SLX4 independent mechanism for MUS81 recruitment (Fig. 2E) while indicating that MUS81 may bind elsewhere in the N-terminal region of TOPBP1 (Fig. 3B).

To address the role of SLX4 pThr1260 in mediating interaction with TOPBP1 and components of the SMX complex, we generated a Thr1260Ala eGFP-SLX4 mutant expression construct and performed co-immunoprecipitations from mitotic synchronised cells after transient expression. Strikingly, mutating SLX4 Thr1260 to alanine abolished binding with TOPBP1 but had no discernible effect on the interaction with MUS81 (Fig. 4B), providing further evidence for a TOPBP1-independent interaction mechanism for these proteins.

Therefore, we conclude that the TOPBP1-SLX4 interaction involves direct binding of phosphorylated SLX4-Thr1260 to TOPBP1 BRCTs 1 and 2. In addition, we establish that MUS81 binds to the N-terminal BRCT module of TOPBP1 (Fig. 3B, 3C and 4A), suggesting a stepwise assembly of the SMX complex *via* interactions with distinct N-terminal BRCT domains of TOPBP1.

### CDK1-dependent phosphorylation of threonine 1260 of SLX4 promotes its interaction with TOPBP1

The mitotic kinase CDK1 is maximally active at the G2-M transition ^49–51^ and SLX4 Thr1260 resides within a predicted minimal CDK consensus site ^28^. Therefore, we hypothesised that the TOPBP1-SLX4 interaction is likely regulated by CDK1. To test this, we raised a custom, phospho-specific antibody against a peptide encompassing phosphorylated SLX4-Thr1260. Consistent with mitotic-specific phosphorylation at SLX4-Thr1260, the antibody preferentially detected phosphorylation in cells isolated by mitotic shake off, which was abolished by phosphatase treatment, indicating its phospho-specificity (Fig. S4A). This phosphorylation was markedly reduced when cells were acutely treated with a CDK1 inhibitor (CDK1i) after mitotic synchronisation, indicating that CDK1 is required for SLX4 phosphorylation (Fig. 4C and D).

To ascertain the functional consequences of this phosphorylation, we examined the interaction between eGFP-SLX4 and TOPBP1, MUS81, and ERCC1 after acute mitotic inhibition of CDK1 (Fig. 4C). Acute mitotic inhibition of CDK1 reduced SLX4 phosphorylation of Thr1260 and binding of TOPBP1 but did not affect its interaction with MUS81 or ERCC1 (Fig. 4C). Similarly, CDK1i treatment hindered the interaction between eGFP-TOPBP1 and SLX4 in reciprocal immunoprecipitation experiments but did not affect the interaction between TOPBP1 and components of the BTR complex (Fig. 4D). Notably, CDK1i treatment also disrupted the interaction between TOPBP1 and MUS81, suggesting that this specific interaction may also be phosphorylation-dependent and CDK1-regulated (Fig. 4D). Consistently, we observed a marked reduction in SLX4 Thr1260 phosphorylation following acute CDK1 inhibition, in an eGFP-TOPBP1 Co-IP (Fig. 4D).

To corroborate the direct phosphorylation of SLX4 by CDK1, we reconstituted the CDK1-SLX4 phosphorylation cascade *in vitro* using purified recombinant CDK1-Cyclin B-CKS1 complex (Fig. S4B), GST-TOPBP1 BRCTs 0-1-2, and a biotinylated-SLX4 Thr1260 peptide in the presence of ATP and Mg^2+^. As such, only streptavidin pull-down of the biotinylated-SLX4 Thr1260 peptide in the presence of TOPBP1 BRCTs 0-2, ATP, Mg^2+^ and recombinant CDK1-Cyclin B-CKS1 facilitated the detection of SLX4 Thr1260 peptide-TOPBP1 BRCT 0-1-2 interaction, comparable to incubation of a pThr1260 peptide with TOPBP1 BRCT 0-1-2 (Fig 4E). This experiment confirmed a direct and specific phosphorylation of SLX4 by CDK1-Cyclin B-CKS1, that may facilitate the mitotic SLX4-TOPBP1 interaction (Fig. 4E).

Therefore, mitotic phosphorylation of SLX4 by CDK1 regulates its interaction with TOPBP1 and this interaction is mediated by the direct binding of phospho-Thr1260 to the BRCT 0-1-2 domains of TOPBP1.

### Threonine 1260 of SLX4 is required for recruitment of components of the SMX complex to chromatin during mitosis and to promote genome stability

To establish the functional role of the mitotic CDK1 driven TOPBP1-SLX4 interaction in SLX4 recruitment, we generated SLX4^−/−^ cells by CRISPR/Cas9 mediated genome editing and complemented them with either inducible wild-type (WT) eGFP-SLX4 or the Thr1260Ala mutant that cannot be phosphorylated to mediate interaction with TOPBP1. Subsequently, we induced expression by addition of doxycycline (Fig. S5A), then treated these cells without or with aphidicolin, to induce replication stress. As TOPBP1 is complexed with CIP2A in mitosis we assessed SLX4 colocalisation with CIP2A on prometaphase chromatin (Fig. 5A).

**Fig. 5:**
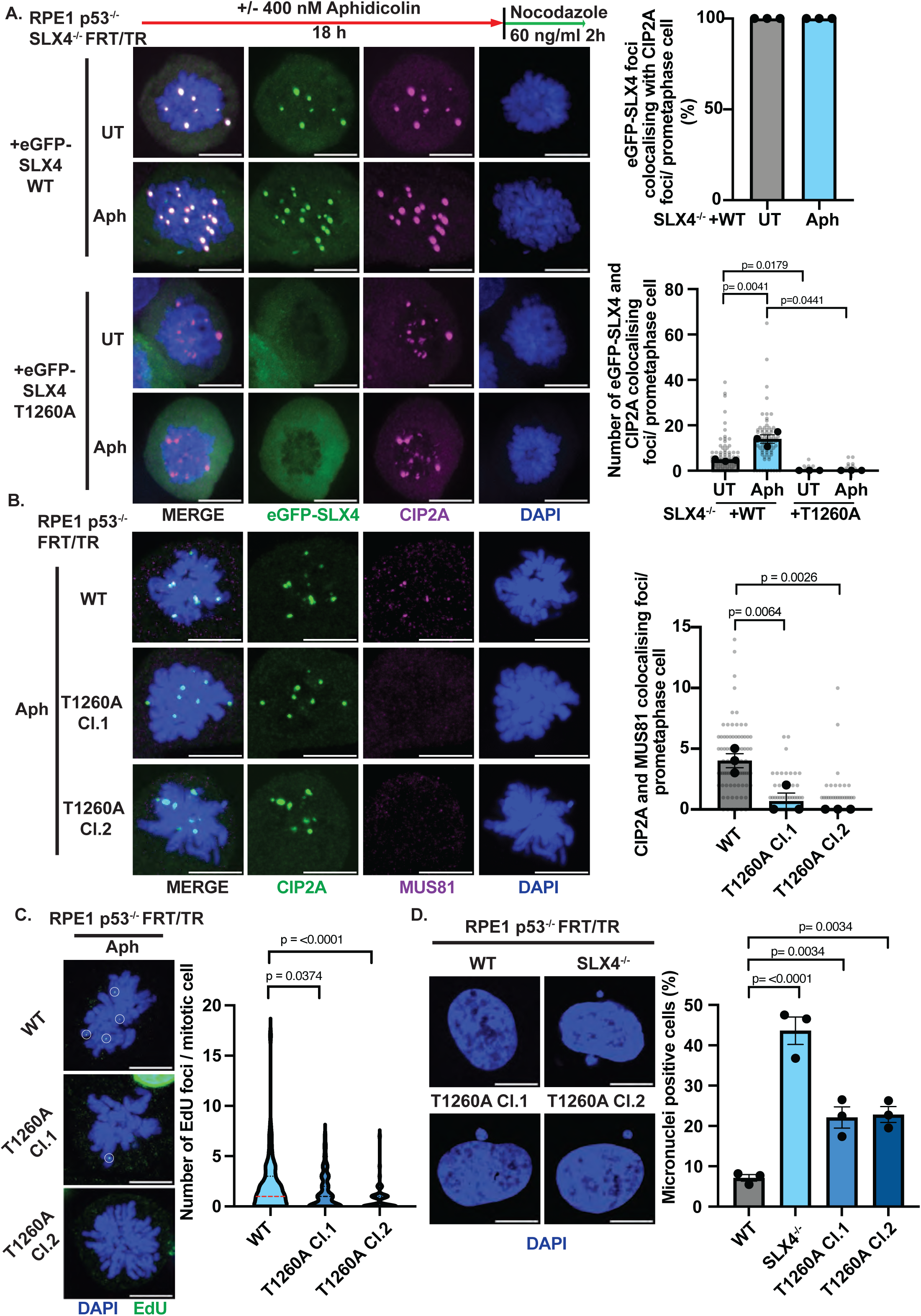
Mitotic localisation of the SMX complex components are CIP2A-TOPBP1 dependent and facilitates unscheduled DNA synthesis to safeguard genome stability. (A) Representative images and (top right panel) a bar plot showing the mean percentage of eGFP-SLX4 WT foci colocalising with CIP2A per RPE1 p53^−/−^ SLX4^−/−^ FRT/TR +eGFP-SLX4 WT/ T1260A prometaphase cell after induction with 10 ng/ml doxycycline for 24 hours followed by treatment with or without 400 nM/ 18 h aphidicolin followed by synchronisation with 60 ng/ml nocodazole for 2 hours (UT: n=85, Aph: n=76). Black dots represent means from each experiment and bars represent mean of three-experiments; (bottom right panel) a dot plot showing the number of eGFP-SLX4 WT or eGFP-SLX4 T1260A foci colocalising with CIP2A foci per prometaphase cell after treatment as above (eGFP-SLX4 WT (UT: n=85, Aph: n=76);) and eGFP-SLX4 T1260A (UT: n=70, Aph: n=76) from three independent experiments, statistical significance was determined by two-way ANOVA). Grey dots represent individual measurements, black dots represent the medians of each experiment and bars show the mean with error bars showing S.E.M. (B) Representative images and a dot plot showing CIP2A and MUS81 colocalising foci in RPE1 p53^−/−^ FRT/TR WT and SLX4 T1260A prometaphase cells treated with 400 nM aphidicolin followed by synchronisation with 60 ng/ml nocodazole for 2 hours (Parental WT: n=80, T1260A Cl.1: n= 76, T1260A Cl.2 n=85, from three independent experiments, statistical significance was determined by one-way ANOVA test with Dunnet’s post-hoc test). Grey dots represent individual measurements, black dots represent the medians of each experiment and bars show the mean with error bars showing S.E.M. (C) Representative images and violin plot of the number of EdU foci in RPE1 p53^−/−^ FRT/TR WT and T1260A prometaphase cells (Parental WT: n=90, T1260A Cl.1: n= 87, T1260A Cl.2 n=82, from three independent experiments, statistical significance was determined by Mann-Whitney U test). (D) Representative images and bar plot of the mean of the percentage of micronuclei-positive RPE1 p53^−/−^ FRT/TR WT, SLX4^-/-^and SLX4 T1260A cells (Parental WT n=336, SLX4^−/−^ n=346, T1260A Cl.1: n= 384, T1260A Cl.2 n=377 from three independent experiments, statistical significance was determined by one-way ANOVA with Dunnet’s post-hoc test), black dots represent the mean of each experiment and bars show the mean with error bars showing S.E.M. Scale bars in all A-D are equivalent to 10 µm.

Subsequently, we observed that WT SLX4 was recruited to prometaphase chromatin in response to replicative stress and colocalised with CIP2A (Fig. 5A). In contrast to WT SLX4, the Thr1260Ala mutant displayed total abrogation of chromatin recruitment in mitosis (Fig. 5A). Importantly, the Thr1260Ala SLX4 mutant did not hinder CIP2A recruitment to prometaphase chromatin (Fig. S5B) and displayed comparable competency to WT in its ability to form foci in interphase cells (Fig. S5C).

To assess the functional role of the TOPBP1-SLX4 interaction in mitotic recruitment of MUS81 and ERCC1, we used a CRISPR-Cas9 targeting strategy to generate cells harbouring the SLX4 Thr1260Ala mutation by knock-in at the endogenous locus (Fig S5D, S6A and B). Following the induction of replication stress with aphidicolin, we examined the colocalisation of CIP2A with MUS81 and ERCC1 (Fig. 5B and S5E). In two indepdent clones of Thr1260Ala knock-in mutant cells, the co-localisation of MUS81 and ERCC1 with CIP2A was significantly disrupted, in stark contrast to their robust colocalisation in WT cells (Fig. 5B and S5E).

Given that SLX4 and MUS81 are required for MiDAS we hypothesised that the TOPBP1-SLX4 interaction may be required to facilitate this mitotic DNA repair mechanism ^11, 12, 52^. Therefore, we treated WT and Thr1260Ala mutant cells with aphidicolin and monitored EdU incorporation in prometaphase cells (Fig. 5C). While WT cells displayed robust EdU incorporation, it was significantly reduced in two independently derived Thr1260Ala mutant clones, indicating an impairment of MiDAS in SLX4 Thr1260Ala cells (Fig. 5C). Consistent with this, cells expressing the SLX4 Thr1260Ala mutant, displayed elevated levels of genome instability, as indicated by micronuclei formation (Fig. 5D). SLX4^−/−^ cells displayed a more pronounced phenotype than the Thr1260Ala knock-in clones (Fig. 5D), reflecting the mitosis-specific function of Thr1260 phosphorylation as opposed to the broader role of SLX4 in DNA repair during both interphase and mitosis.

Collectively, these findings determine that the functional role of the TOPBP1-SLX4 interaction in human cells is to facilitate mitotic recruitment of the SMX complex components, SLX4, MUS81, and ERCC1, to sites of DNA damage marked by CIP2A-TOPBP1 during mitosis, to drive MiDAS and safeguard genome stability.

### CIP2A regulates SLX4 and Polθ recruitment to orchestrate mitotic DNA repair

The role of CIP2A in the regulation of DNA repair activity is unknown. We hypothesised that CIP2A acts upstream of SLX4 given that SLX4 depletion did not alter TOPBP1 recruitment and the Thr1260Ala mutation also not impair mitotic CIP2A localisation (Fig. 2D, Fig. 5A, B and Fig. S5E). To address this, we analysed the mitotic-specific recruitment of eGFP-SLX4 WT and its colocalisation with TOPBP1 in prometaphase cells where CIP2A was present or depleted using validated siRNA ^1, 13^ (Fig. S7A). Consistent with our hypothesis, we observed that SLX4 colocalised with TOPBP1 in mitosis (Fig. 6A and B), and a near-complete absence of SLX4 and TOPBP1 colocalising foci on prometaphase chromatin in CIP2A-depleted cells (Fig. 6C). As expected, we also observed a reduction in TOPBP1 foci formation on prometaphase chromatin in CIP2A depleted cells (Fig. S7B).

**Fig. 6:**
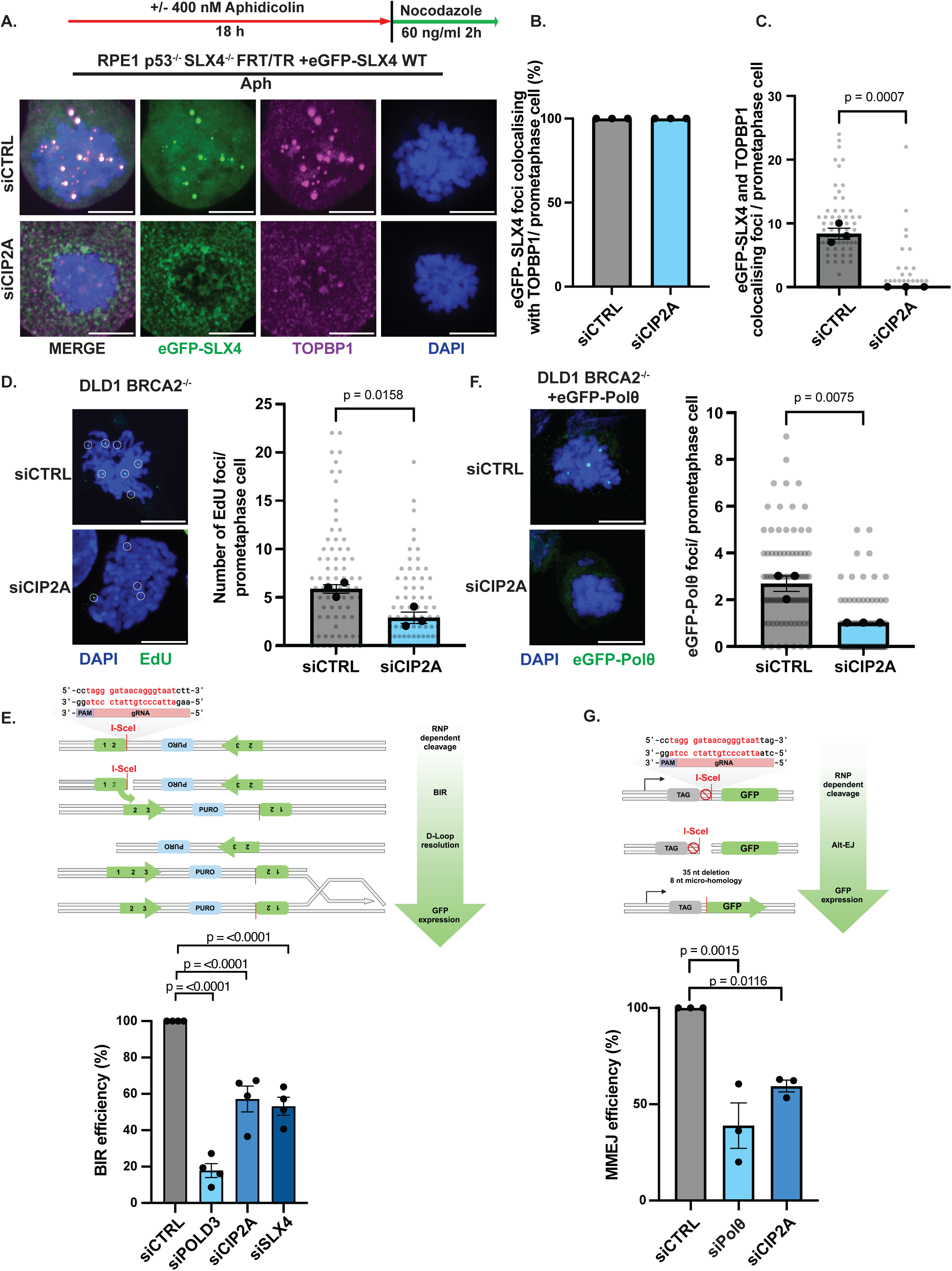
CIP2A is required for the regulation of redundant mitotic repair pathways. (A) Representative images and (B) a bar plot showing the percentage of eGFP-SLX4 foci colocalising with TOPBP1 in RPE1 p53^−/−^FRT/TR SLX4^−/−^ prometaphase cells and (C) number of eGFP-SLX4 and TOPBP1 colocalising foci in RPE1 p53^−/−^FRT/TR prometaphase cells induction with 10 ng/ml doxycycline for 24 hours, 48 hours after siRNA treatment. (B and C) Cells were treated with siCTRL or siCIP2A (siCTRL: n=73, siCIP2A: n=73, statistical significance was determined by two tailed unpaired t-test). (D) Representative images and dot plot of EdU foci in prometaphase cells DLD1 BRCA2^−/−^cells (siCTRL: n=75, siCIP2A: n= 74 from three independent experiments, statistical significance was determined by two tailed unpaired t test). (E) Reporter system schematic (created with BioRender.com) and bar plot of BIR dependent repair efficiency following sgRNP-CAS9 facilitated cleavage of an I-SceI site in the pBIR-GFP BIR reporter U2OS cell line, following treatment with siCTRL, siPOLD3, siCIP2A or siSLX4. Measurements are from four independent experiments. (F) Representative images and dot plot of eGFP-Polθ foci in prometaphase DLD1 BRCA2^−/−^ eGFP-Polθ cells after induction with 100 ng/ml doxyclycline for 24 hours, 48 hours after siRNA treatment. (siCTRL: n=86, siCIP2A: n= 83 from three independent experiments, statistical significance was determined by two tailed unpaired t-test). In C, D and F grey dots represent individual measurements; black dots represent medians of each experiment and bars the mean with S.E.M displayed. Scale bars equivalent to 10 µm. (G) Reporter system schematic (created with BioRender.com) and bar plot of MMEJ repair efficiency following sgRNP-CAS9 generated cleavage of an I-SceI site in the EJ2 reporter U2OS cell lines following treatment with siCTRL, siPolθ or siCIP2A. Measurements are from three independent experiments. Statistical significance in E and G was determined by one-way ANOVA with Dunnet’s post-hoc test and the mean is presented as bars with S.E.M.

As BRCA2 deficient cells demonstrate elevated reliance on MiDAS (Fig. S7C) ^53^, we analysed MiDAS by measuring EdU incorporation in mitotic DLD1 BRCA2^−/−^ cells depleted of CIP2A by siRNA (Fig. 6D and S7D). We observed a significant reduction in EdU incorporation in CIP2A-depleted cells (Fig. 6D), highlighting a pivotal role for CIP2A in facilitating MiDAS. Since MiDAS operates through a break-induced replication (BIR)-like mechanism ^19^, we sought to strengthen this observation by investigating the role of CIP2A in BIR-dependent DNA repair utilising a BIR reporter (Fig. 6E) ^54,55^. As expected, depletion of POLD3 or SLX4, known drivers of MiDAS, significantly impaired efficiency of DSB repair when compared to siCTRL treated cells (Fig. 6E and fig. S7E). In line with our prediction, CIP2A depletion also significantly reduced BIR efficiency (Fig. 6E and fig. S7E), providing direct evidence that CIP2A is critical for the repair of DSBs by BIR.

Recently Gelot *et al.* demonstrated that Polθ phosphorylation by PLK1 facilitates its interaction with TOPBP1, thereby driving mitotic MMEJ-dependent repair of DSBs ^5^. Given the mutual dependency between CIP2A and TOPBP1 for chromatin recruitment during mitosis ^1, 13^, we hypothesised that CIP2A may also regulate mitotic MMEJ by facilitating the mitotic recruitment of Polθ to mitotic chromatin. To test this hypothesis, we generated DLD1 BRCA2^−/−^ cells that express eGFP-tagged Polθ in an inducible manner and analysed its recruitment to mitotic chromatin following CIP2A depletion (Fig. S7F). In siCTRL treated cells, eGFP-Polθ was recruited to prometaphase chromatin; however, this recruitment was significantly reduced in CIP2A-depleted cells (Fig. 6F and fig. S7F). To directly assess the role of CIP2A in MMEJ, we used an additional U2OS cell line expressing a MMEJ reporter construct (Fig. 6G) ^56^. Depletion of Polθ in this system led to a significant defect in MMEJ efficiency, validating our approach. Strikingly, we observed that CIP2A depletion also led to a marked impairment in MMEJ (Fig. 6G and S7G), which is consistent with the observed defective recruitment of Polθ upon CIP2A depletion (Fig. 6F). Together these findings demonstrate a critical role for CIP2A in facilitating not only BIR-like MiDAS but also Polθ-mediated MMEJ during mitosis.

Next, we examined the impact of CIP2A loss (Fig. S7H) in comparison to the combined disruption of the TOPBP1-SLX4 interaction (required for MiDAS) and Polθ (required for MMEJ) inhibition on genome stability in response to replication stress induced by aphidicolin. We observed an increase in micronuclei formation in response to Polθ inhibition, although this increase was less pronounced than that observed in SLX4 Thr1260Ala mutant cells (Fig. 7A). Notably, inhibition of Polθ (required for MMEJ) in SLX4 Thr1260Ala mutant cells (required for MiDAS) led to an even greater increase in micronuclei formation, indicating an additive effect. Consistent with the role of CIP2A in facilitating both MMEJ and MiDAS repair in mitosis its depletion in WT cells showed similar levels of micronuclei formation to that observed in SLX4 Thr1260Ala cells treated with Polθ (Fig. 7A and S7H). Furthermore, analysis of cellular proliferation in the presence of aphidicolin revealed that SLX4 Thr1260Ala mutant knock-in cells displayed a defect in proliferation that was additive when incubated in the presence of a Polθ inhibitor (Fig.7B). Finaly, depletion of SLX4 in combination with Polθi resulted in a broadly similar loss of proliferative capacity compared to single depletion of CIP2A alone in BRCA2 deficient DLD1 cells (Fig. 7C). Consistent with BRCA1/2 deficient cells being reliant on the TOPBP1-SLX4 interaction to facilitate MiDAS and safeguard genome stability, we also observed that induction of expression of a minimal SLX4 Thr1260 containing fragment in DLD1 BRCA2^−/−^ (Fig. S8A) or SUM149PT BRCA1^−/−^ (Fig. S8B) cells was sufficient to impair cellular proliferation.

**Fig. 7.**
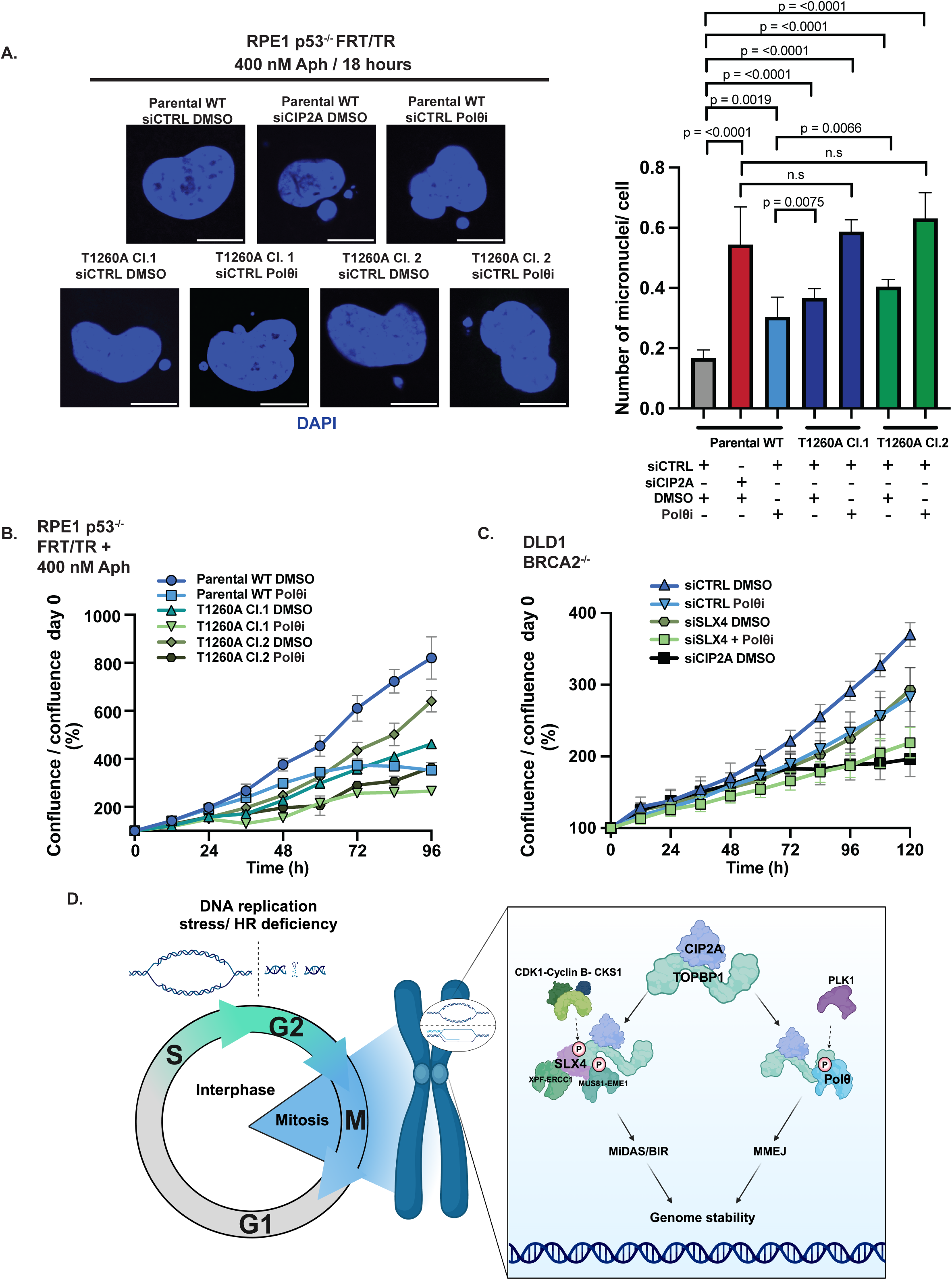
CIP2A dependent orchestration of DNA repair is required for genome stability and cellular proliferation in the absence of BRCA2. (A) Representative images and bar plot illustrating the number of micronuclei per RPE1 p53^−/−^ FRT/TR WT or SLX4 T1260A cells. Cells were treated with siCTRL or siCIP2A, 400 nM aphidicolin and with either DMSO or 5 µM ART558 (Polθi) for 18 hours (Parental WT (siCTRL + DMSO: n =343, siCIP2A + DMSO: n=394, siCTRL + Polθi: n=372), SLX4 T1260A Cl.1 (siCTRL + DMSO: n= 385 and siCTRL + Polθi: n =341) or SLX4 T1260A Cl.2 (siCTRL + DMSO: n= 399 and siCTRL + Polθi: n =378) from three independent experiments). Statistical significance was determined using by the Mann-Whitney test, and the mean of three experiments is presented as bars with S.E.M. (B) Proliferation analysis of RPE1 p53^−/−^ FRT/TR WT and T1260A cells, performed using the Incucyte live cell analysis system. Cells were treated with or without 5 µM ART558 (Polθi) in the presence of 400 nM aphidicolin. The mean confluence divided by confluence at day 0, from three independent experiments, is displayed with S.E.M. Scale bars equivalent to 10 µm. (C) Proliferation analysis of DLD1 BRCA2^−/−^ cells using the Incucyte live cell analysis system. Cells were treated with siCTRL, siSLX4, siCIP2A in the presence of DMSO or 5 µM ART558 (Polθi). The mean confluence divided by confluence at day 0, from three independent experiments, is displayed with S.E.M. (D) Model: Cells enter mitosis with DNA damage, under replicated DNA or recombination intermediates where the CIP2A-TOPBP1 complex is recruited. CDK1 phosphorylation of SLX4 at T1260 regulates interaction with TOPBP1 BRCT 1/2 facilitating SMX component recruitment and MiDAS/BIR to safeguard genome stability. PLK1 phosphorylation of POLQ in mitosis facilitates TOPBP1 interaction and MMEJ. Loss of CIP2A impairs both pathways and leads to simultaneous deficiency in mitotic MiDAS/BIR and MMEJ promoting genome instability (created with BioRender.com).

We conclude that CIP2A is essential for facilitating mitotic DNA double-strand break repair through multiple mitotic DNA repair pathways, including coordination of BIR-like processes through (TOPBP1-SLX4) and MMEJ through Polθ. This work identifies its critical role in maintaining genome stability during mitosis and provides a mechanistic basis for its synthetic lethality in BRCA1/2-deficient cells (Fig. 7D).

## Discussion

Recent studies have underscored the importance of mitosis as a critical phase for the maintenance of genome stability, with the CIP2A-TOPBP1 axis identified to have a pivotal role in these processes, speculated to be through promoting chromatin tethering ^8, 32, 33^. Yet direct evidence for role of the CIP2A complex in regulation of mitotic DNA repair pathways remains absent. In addition, the mitotic TOPBP1-Polθ interaction facilitated by PLK1-dependent phosphorylation, has been shown to be essential for the recruitment of Polθ to mitotic chromosomes to facilitate MMEJ repair^5^. Notably, elegant work from several laboratories has demonstrated that the CIP2A-TOPBP1 and TOPBP1-Polθ interactions are crucial for the survival of BRCA1/2-deficient cancers, with mitotic-specific disruption of CIP2A-TOPBP1 or TOPBP1-Polθ being sufficient to inhibit cell growth in BRCA2-deficient cells but not in wild-type cells ^1, 5^. However, the underlying mechanism by which CIP2A loss leads to synthetic lethality in BRCA1/2-deficient cells remains unclear, particularly since chromosome tethering alone does not fully explain this phenotype, given that MDC1, which also plays a role in mitotic chromosome tethering, does not exhibit synthetic lethality with BRCA1/2 loss^32–35^.

In this study, we address this critical question by determining that the CIP2A-TOPBP1 complex plays a critical role in facilitating mitotic DSB repair pathway choice, through the control of mitotic SMX complex and Polθ-dependent recruitment. As such, we demonstrate that the mitotic kinase CDK1 directly orchestrates the assembly of the SMX complex components SLX4, MUS81 and ERCC1 at sites marked by the CIP2A-TOPBP1 complex in mitosis. This is achieved by phosphorylation of SLX4 at Thr1260 which is critical for interaction between SLX4 and BRCT1/2 domains of TOPBP1. This phosphorylation event serves as a trigger, which subsequently drives DSB repair via a break-induced replication (BIR)-like MiDAS process to safeguard genome stability. Since phosphorylation of SLX4 at Thr1260 enhances its interaction with TOPBP1 in yeast^28, 29^, our work identifies an evolutionary highly conserved mechanism mediating DSB via BIR/MiDAS during mitosis. Furthermore, we demonstrate that targeting this interaction via expression of a minimal SLX4 fragment in BRCA1/2 deficient cancer cells, that are characterised to exhibit heightened replicative stress, significantly impairs cellular fitness. This mechanism provides rationale for the synthetic lethality observed between SLX4 or MUS81 loss in BRCA2 deficiency^53^.

Our findings indicate that CIP2A is not only crucial for the mitotic-specific recruitment of TOPBP1 and the SMX complex but also for the recruitment of Polθ to mitotic chromatin, thus playing a central role in orchestrating both mitotic BIR and MMEJ. Accordingly, the combination of Polθ inhibition with the SLX4 Thr1260Ala mutation leads to a significantly higher level of genome instability and impaired cellular fitness compared to either mutation alone. This defines SLX4 and Polθ, as downstream effectors of the CIP2A-TOPBP1 complex that act non-epistatically to support genome stability and cellular fitness. Interestingly, and in support of our observations in human cells Carvajal-Garcia *et al*. defined that combined loss of Polθ, SLX4 and BRCA2 hypersensitised *D. melanogaster* cells to ionizing radiation and replication stress induced by camptothecin^57^.

An interesting aspect of our findings is the stepwise assembly of components of the SMX complex and Polθ in mitosis in a CIP2A-TOPBP1 dependent manner, orchestrated through the regulatory control of mitotic kinases CDK1 and PLK1 ^5^. Specifically, CDK1-driven phosphorylation of SLX4 at Thr1260 facilitates its interaction with the BRCT1/2 domains of TOPBP1, promoting the recruitment of the SMX complex components to sites of DNA damage. Concurrently, PLK1-mediated phosphorylation of Polθ was shown to enable the recruitment of Polθ, thus enhancing MMEJ repair activity during mitosis^5^. This bi-modal activation process ensures both spatial control via the TOPBP1-CIP2A complex and temporal control via mitotic kinases CDK1-PLK1, thereby tightly regulating the engagement of mitotic DNA repair pathways. This coordinated assembly and regulation of repair complexes underscores the multifaceted role of CIP2A-TOPBP1 as a central hub in maintaining genomic stability during cell division.

Intriguingly, CIP2A depletion in MEFs does not affect telomere fusions in the absence of TRF2 ^36^ which are dependent on MMEJ^10^. This suggests that either CIP2A’s role in promoting MMEJ may differ in mouse cells, or that MMEJ-dependent telomeric fusions occur largely outside of mitosis i.e. in late S/G2 phase. In addition, MMEJ driven telomere fusions may involve an alternative mechanism of Polθ recruitment and regulation—potentially mediated by RHNO1, as recently demonstrated, rather than CIP2A^10^. Finally, while our study provides evidence for the critical role of CIP2A in coordinating MiDAS and MMEJ DNA repair activities during mitosis, we do not rule out the possibility of CIP2A-TOPBP1-independent MMEJ during S phase or even mitosis ^10, 58, 59^.

This work offers strong rationale for the targeting of these pathways in tumours that are BRCA1/2-deficient, that exhibit heightened endogenous replicative stress or in combination with established anti-cancer treatment modalities that target DNA replication. Future research aimed at unravelling the precise dynamics and full complement of mitotic DNA repair mechanisms, will be crucial for unlocking the untapped therapeutic potential associated with targeting these pathways.

## Supporting information

Supplementary figures and methodology

## Acknowledgements

We thank D. Cortez (Vanderbilt University) for HCT116-osTIR1-TOPBP1-mAID-Clover cells, C. Lord for DLD1 and DLD1 BRCA2^−/−^ cells and SUM149PT cells, and S. Jackson for RPE1 p53^−/−^ FRT/TR cells. Thanos D. Halazonetis for providing U2OS-pBIR reporter cells and Jeremy Stark for providing U2OS-EJ2 reporter cells.

The ICR Light Microscopy Core facility for assistance with use of microscopes and image analysis ICR (K. Betteridge, R. Scrimgeour and Q. Lai) H. Ale and M. Guelbert) for assistance with microscopy and flow cytometry cell sorting. The ICR Flow Cytometry Facility for assistance with cell sorting and cell cycle analysis (H.Ale, Y. Semochkina and B. Satam). We also thank G. Benstead-Hume and S. Haider for bioinformatical analysis. Schematic graphics were created using BioRender.com. We thank C. Zerhuit, G. Coster and J. Pines for feedback on the manuscript ahead of submission.

This work was funded by an MRC Research Grant to W.N and P.M (MRC295X) and a BBSRC Discovery fellowship to P.M (BB/T009608/1). Work in W.N’s lab was also supported by a CR-UK Programme grant [A24881]. J.M lab and J.V were funded by a Cancer Research UK Senior Cancer Research Fellowship to J.M. (RCCSCF-Nov22/100001). Funding supporting Z.K. and J.S.C. was provided by Wellcome Trust [223745/Z/21/Z] and from ICR core funding. Work carried out by M.D, A.O and L.P was supported by a Cancer Research UK Programme Grant (C302/A24386) and a Royal Society Research Grant RG\R2\232017. Work in the J.D lab was supported by Cancer Research UK (C7905/A25715).

## Author contributions

P.M. and W.N conceived the project. P.M. and W.N designed experiments with contribution from M.D, J.N, Z. K, J.C, L.P and A.O. P.M. performed most of the experiments. J.N., M.D., Z.K, J. V, A.O, N.J, M.L, A.K, K.L, and A.M, contributed to specific experiments. Z.K and J.C assisted with proteomics experimental design and analysis. M.D, A.O, and L.P assisted with *in vitro* experimental design, protein purification, fluorescence polarization and CDK1 phosphorylation reconstitution experiments. J.V and J. M assisted with CRISPR knock in design and execution. W.N and P.M wrote the manuscript, and all authors reviewed the manuscript ahead of submission.

