## Supplementary figures and methodology for "The mitotic CIP2A-TOPBP1 axis facilitates mitotic pathway choice between MiDAS and MMEJ"

### Supplementary figure legends

**Fig.S1:**

(A) Representative histogram plots of flow-cytometry analysis of propidium iodide stained HEK293TN asynchronous, S-phase synchronised and M-phase synchronised cells, as carried out in Fig 1 for Co-IP and mass-spectrometry analysis.

(B) Representative images and treatment schematic of HCT116-TOPBP1-mAID-Clover cells treated with or without 500 μM IAA as indicated. (B) Bar plots of anaphase abnormalities upon acute TOPBP1 loss. (Left panel) (C) Percentage of anaphases with DAPI marked chromatin bridges. (Middle panel) Percentage of anaphase cells with DAPI marked DNA laggards, (-: n= 111, +: =125 from four independent experiments). (Right panel) Percentage of anaphases with PICH marked ultra-fine anaphase bridges (-: n= 40, +: =47 from two independent experiments). Statistical significance in C was determined by two tailed unpaired t-tests. Scale bars equivalent to 10 μM. (D) Western blot showing degredation analysis of endogenously mAID-Clover tagged TOPBP1 in HCT116-ostir1 cells with or without treatment with 500 μM IAA as indicated. (E) Western blot analysis of BLM, ERCC1, MUS81 and SLX4 interactions with eGFP-TOPBP1 in GFP-TRAP Co-IPs from HEK293TN cells transiently transfected with a eGFP-TOPBP1 WT expression construct followed with or without mitotic synchronisation by 100 ng/ml nocodazole for 16 hours. 1% of input was used for analysis by western blot of input lysate.

**Fig.S2:**

(A) Representative images and dot plot of TOPBP1 foci in DLD1 and DLD1 BRCA2$^{-/-}$ prometaphase cells (WT: n=74, BRCA2$^{-/-}$: n=73 from three independent experiments, statistical significance was determined by two tailed unpaired t-test). Grey dots represent individual measurements, black dots indicate medians from individual experiments and bars represent the mean with S.E.M displayed. (B) Representative images and violin plot of MUS81 and TOPBP1 colocalising foci in RPE1 p53$^{-/-}$ FRT/TR prometaphase cells treated with or without 400 nM aphidicolin for 18 hours followed by 2 hour nocodazole treatment (UT:n= 96, Aph :n= 134, from two independent experiments). (C) Representative images and dot plot of ERCC1 and TOPBP1 colocalising foci in RPE1 p53$^{-/-}$ FRT/TR prometaphase cells treated with or without 400 nM aphidicolin for 18 hours followed by 2 hour nocodazole treatment (UT:n= 57, Aph :n= 47, from two independent experiments). Statistical significance in B and C was determined by Mann Whitney test, red line represents median. (D). Representative images and dot plot of TOPBP1-mAID-clover foci in HCT116-TOPBP1-mAID-Clover cells treated with or without IAA for 2 h. (-: n=86, +:n =88 from three independent experiments, statistical significance was determined by two tailed unpaired t-test). Grey dots represent individual measurements, black dots indicate medians from individual experiments and bars represent the mean with S.E.M displayed. Scale bars in A, B , C and D are equivalent to 10 μm.

(E) Western blot analysis of DLD1 and DLD1 BRCA2$^{-/-}$ cells treated with siCTRL or siTOPBP1.

(F) Western blot analysis of of DLD1 and DLD1 BRCA2$^{-/-}$ cells treated with siCTRL or siSLX4.

**Fig.S3:**

(A) Domain organisation of eGFP-TOPBP1proteins analysed in and used in Fig. 3A. (B) SLX4 domain organisation and location of candidate TOPBP1 BRCT 1 interacting SLX4 T1476 residue and fluorescence polarisation analysis of fluorescein labelled SLX4 pT1476 containing peptide in the presence of recombinant tandem BRCT containing TOPBP1 fragments.

**Fig.S4**
(A) Western blot analysis of pT1260 SLX4 and total SLX4 in asynchronous, interphasic or mitotic cells isolated by mitotic shake off treated with or without shrimp intestinal phosphatase combined with $\lambda$ phosphatase and shrimp intestinal phosphatase (dashed line indicates vertical separation to facilitate indepdent incubation of membrane with phosphatases (see methods)). (B) SDS-PAGE showing purified recombinant CDK1-Cyclin B-CKS1 complex fractions E2-E4 were pooled for use in assays.

**Fig.S5**

(A) Western blot analysis of eGFP-SLX4 expression in RPE1 p53$^{-/-}$ SLX4$^{-/-}$ FRT/TR + eGFP-SLX4 WT and eGFP-SLX4 T1260A cells treated with 10 ng/ml doxycycline for 24 hours as treated in advance of all experiments using cell lines. (B) Representative images and dot plot of CIP2A foci in RPE1 p53$^{-/-}$ SLX4$^{-/-}$ FRT/TR prometaphase cells treated with or without 400 nM aphidicolin for 18 hours followed by 60 ng/ml nocodazole synchronisation (+eGFP-SLX4 WT (UT: n=85, Aph: n=76) +eGFP-SLX4 T1260A (UT: n=70, Aph: n=76) from three independent experiments, statistical significance was determined using two way ANOVA). Grey dots represent individual measurements, black dots represent medians from individual experiments and bars show the mean with S.E.M displayed. (C) Representative images and dot plot of eGFP-SLX4 or eGFP-SLX4 T1260A foci in interphasic RPE1 p53$^{-/-}$ SLX4$^{-/-}$ FRT/TR cells (+eGFP-SLX4 WT (UT: n=565, Aph: n=369) +eGFP-SLX4 T1260A (UT: n=639, Aph: n=421) from three independent experiments, statistical significance was determined by two tailed unpaired t-test on median values from individual experiments). Grey dots show individual measurements. (D) Western blot analysis of SLX4 expression in WT, SLX4$^{-/-}$, SLX4 T1260A knock-in clones (Cl.) 1 and 2. (E) Representative images and dot plot of CIP2A and ERCC1 colocalising foci in RPE1 p53$^{-/-}$ FRT/TR WT and SLX4 T1260A prometaphase cells, treated with 400 nM aphidicolin for 18 hours followed by synchronisation with 60 ng/ml nocodazole for 2 hours (Parental WT: n=88, T1260A Cl.1: n= 80 T1260A Cl.2 n=76, from three independent experiments, statistical significance was determined by one way ANOVA with Dunnet's post hoc test). Grey dots represent individual measurements, black dots represent medians from individual experiments and bars show the mean with S.E.M displayed. Scale bars in A, B and E are equivalent to 10 µm.

**Fig.S6**
(A) Schematic of RPE1 p53$^{-/-}$ FRT/TR SLX4 Thr1260Ala knock-in cell line generation via ATP1A coselection (created with BioRender.com). Multiple-sequence alignment of reference sequence with translated native amino acids and guide RNA target sites labelled, below is alignment of SLX4 Thr1260Ala repair template used, containing Thr1260Ala point mutation, silent mutation of guide RNA recognition sites and silent mutation upstream to create a KpnI restriction site (highlighted by red dashed box). Below shows sanger sequencing traces of WT, Thr1260Ala Cl.1 and Cl.2 PCR products amplified from genomic DNA using primers spanning the edited locus.

**Fig.S7**
(A) Western blot analysis of SLX4 and CIP2A in RPE1 p53$^{-/-}$ SLX4$^{-/-}$ FRT/TR cells treated with 10 ng/ml doxycycline for 24 hours to induce eGFP-SLX4 WT expression, treated with siCTRL or siCIP2A. (B) Representative images of dot plot of TOPBP1 foci in RPE1 p53$^{-/-}$ SLX4$^{-/-}$ FRT/TR eGFP-SLX4 WT expressing prometaphase cells treated with siCTRL or siSLX4 and followed by 400 nM aphidicolin for 18 hours followed by synchronisation with 60 ng/ml nocodazole for 2 hours (siCTRL: n=74, siSLX4: n=73 from three independent experiments, statistical significance was determined using two tailed unpaired t-test). (C) Representative images and dot plot of EdU foci in prometaphase DLD1 WT and BRCA2$^{-/-}$ cells (WT: n=52,

BRCA2$^{-/-}$: n= 52, from two independent experiments, statistical significance was determined by two tailed upaired t-test). In A and B grey dots represent individual measurements, black dots the median from each individual experiment and the bars represent the means with S.E.M displayed. In A-B scale bars are equivalent to 10 µm. (D) Western blot analysis of DLD1 BRCA2$^{-/-}$ treated with siCTRL or siCIP2A. (E) Western blot analysis of U2OS pBIR reporter cells treated with siCTRL, siSLX4, siCIP2A or siPOLD3. (F) Western blot analysis of eGFP-Polθ and CIP2A in DLD1 BRCA2$^{-/-}$ cells treated with siCTRL or siCIP2A followed by 100 ng doxycycline for 24 hours to induce eGFP- Polθ expression. (G) Western blot analysis of U2OS EJ2 reporter cells treated with siCTRL, siCIP2A. (H) Western blot analysis of RPE1 p53$^{-/-}$ FRT/TR cells treated with siCTRL or siCIP2A. (I) Western blot analysis of DLD1 and DLD1 BRCA2$^{-/-}$ cells treated with siCTRL, siSLX4, siCIP2A or siSLX4 and siCIP2A.

**Fig.S8**

(A) Proliferation analysis using the Incucyte S3 live cell analysis system of DLD1 BRCA2$^{-/-}$ cells with inducible expression of empty vector (EV) or a SLX4 T1260 fragment, incubated in the presence of 1 µg/ml doxycycline. Data points represent mean of two independent experiments with error bars representing S.E.M. (B) Proliferation analysis using the Incucyte S3 live cell analysis system of SUM149PT BRCA1$^{-/-}$ cells with inducible expression of empty vector (EV) or a SLX4 T1260 fragment, incubated in the presence of 1 µg/ml doxycycline. Data points represent mean of two independent experiments with error bars representing S.E.M.

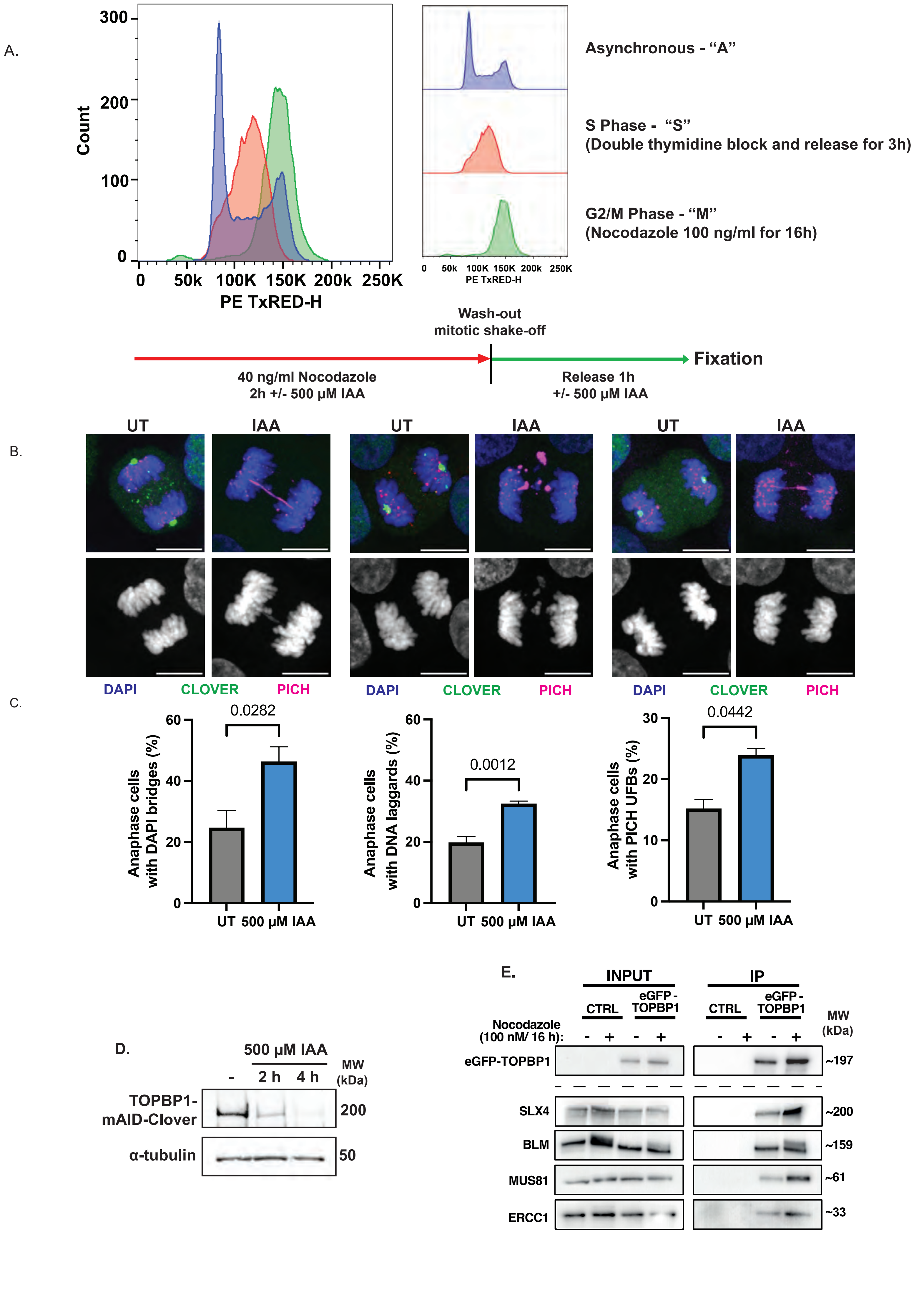
A.
300
200
100
0
Count
0
50k
100K
150K
200k
250K
PE TxRED-H
50k
100K
150K
200k
250K
PE TxRED-H
Asynchronous - "A"
S Phase - "S"
(Double thymidine block and release for 3h)
G2/M Phase - "M"
(Nocodazole 100 ng/ml for 16h)
Wash-out
mitotic shake-off
Fixation
40 ng/ml Nocodazole
2h +/- 500 μM IAA
Release 1h
+/- 500 μM IAA
B.
UT
IAA
UT
IAA
UT
IAA
DAPI
CLOVER
PICH
DAPI
CLOVER
PICH
DAPI
CLOVER
PICH
C.
Anaphase cells
with DAPI bridges (%)
60
40
20
0
0.0282
UT
500 μM IAA
Anaphase cells
with DNA laggards (%)
60
40
20
0
0.0012
UT
500 μM IAA
Anaphase cells
with PICH UFBs (%)
30
20
10
0
0.0442
UT
500 μM IAA
D.
500 μM IAA
-
2 h
4 h
MW
(kDa)
TOPBP1-
mAID-Clover
200
α-tubulin
50
E.
INPUT
IP
CTRL
eGFP -
TOPBP1
CTRL
eGFP -
TOPBP1
MW
(kDa)
Nocodazole
(100 nM/ 16 h):
- + - + - + - +
eGFP-TOPBP1
~197
SLX4
~200
BLM
~159
MUS81
~61
ERCC1
~33

Figure S1

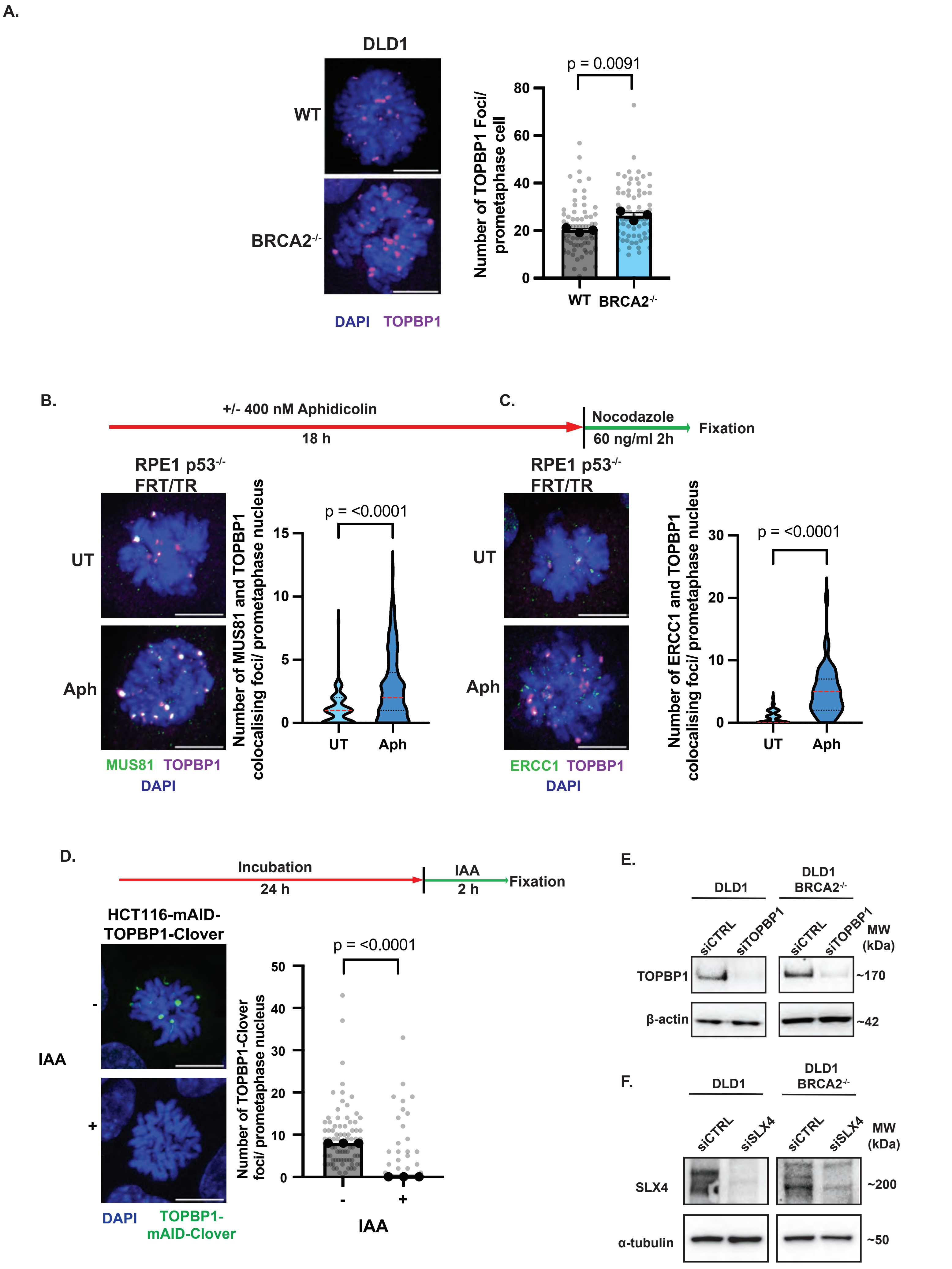

A.
DLD1
WT
BRCA2-/-
DAPI TOPBP1
p = 0.0091
Number of TOPBP1 Foci/ prometaphase cell
80
60
40
20
0
WT
BRCA2-/-
B.
+/- 400 nM Aphidicolin
18 h
C.
Nocodazole
60 ng/ml 2h
Fixation
RPE1 p53-/- FRT/TR
UT
Aph
MUS81 TOPBP1
DAPI
Number of MUS81 and TOPBP1 colocalising foci/ prometaphase nucleus
p = <0.0001
15
10
5
0
UT
Aph
RPE1 p53-/- FRT/TR
UT
Aph
ERCC1 TOPBP1
DAPI
Number of ERCC1 and TOPBP1 colocalising foci/ prometaphase nucleus
p = <0.0001
30
20
10
0
UT
Aph
D.
Incubation
24 h
IAA
2 h
Fixation
HCT116-mAID-TOPBP1-Clover
-
IAA
+
DAPI
TOPBP1-mAID-Clover
p = <0.0001
Number of TOPBP1-Clover foci/ prometaphase nucleus
50
40
30
20
10
0
-
+
IAA
E.
DLD1
DLD1 BRCA2-/-
siCTRL
siTOPBP1
siCTRL
siTOPBP1
MW (kDa)
TOPBP1
~170
β-actin
~42
F.
DLD1
DLD1 BRCA2-/-
siCTRL
siSLX4
siCTRL
siSLX4
MW (kDa)
SLX4
~200
α-tubulin
~50


Figure S2

A.

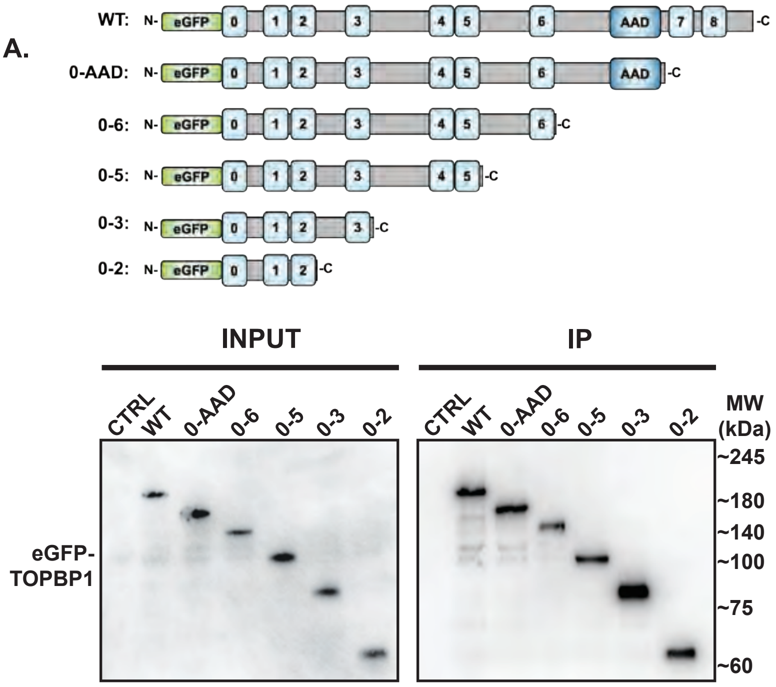

WT: N- eGFP 0 1 2 3 4 5 6 AAD 7 8 -C
0-AAD: N- eGFP 0 1 2 3 4 5 6 AAD -C
0-6: N- eGFP 0 1 2 3 4 5 6 -C
0-5: N- eGFP 0 1 2 3 4 5 -C
0-3: N- eGFP 0 1 2 3 -C
0-2: N- eGFP 0 1 2 -C
INPUT
IP
CTRL WT 0-AAD 0-6 0-5 0-3 0-2
CTRL WT 0-AAD 0-6 0-5 0-3 0-2
MW (kDa)
~245
~180
~140
~100
~75
~60
eGFP-TOPBP1


B.

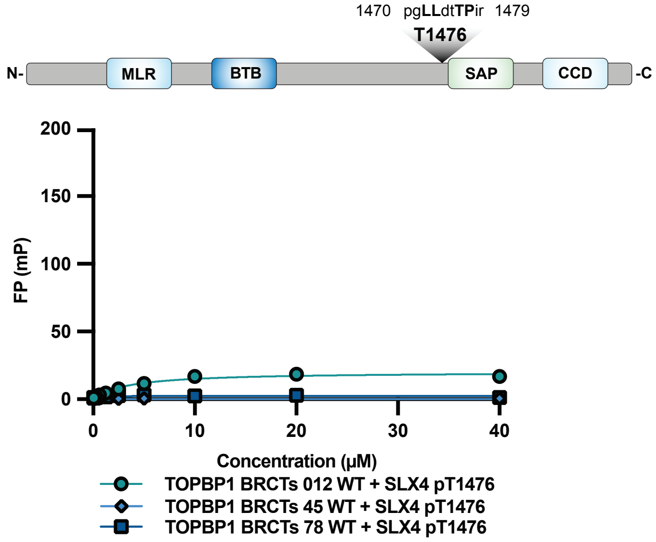

1470 pgLLdtTPir 1479
T1476
N- MLR BTB SAP CCD -C
FP (mP)
200
150
100
50
0
0
10
20
30
40
Concentration (μM)
TOPBP1 BRCTs 012 WT + SLX4 pT1476
TOPBP1 BRCTs 45 WT + SLX4 pT1476
TOPBP1 BRCTs 78 WT + SLX4 pT1476


Figure S3

A.

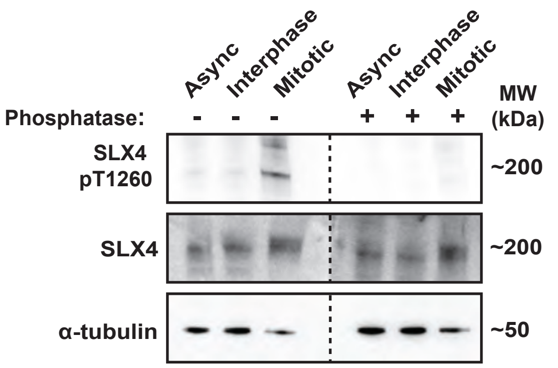
Async
Interphase
Mitotic
Async
Interphase
Mitotic
MW (kDa)
Phosphatase: - - - + + +
SLX4 pT1260
~200
SLX4
~200
α-tubulin
~50

B.

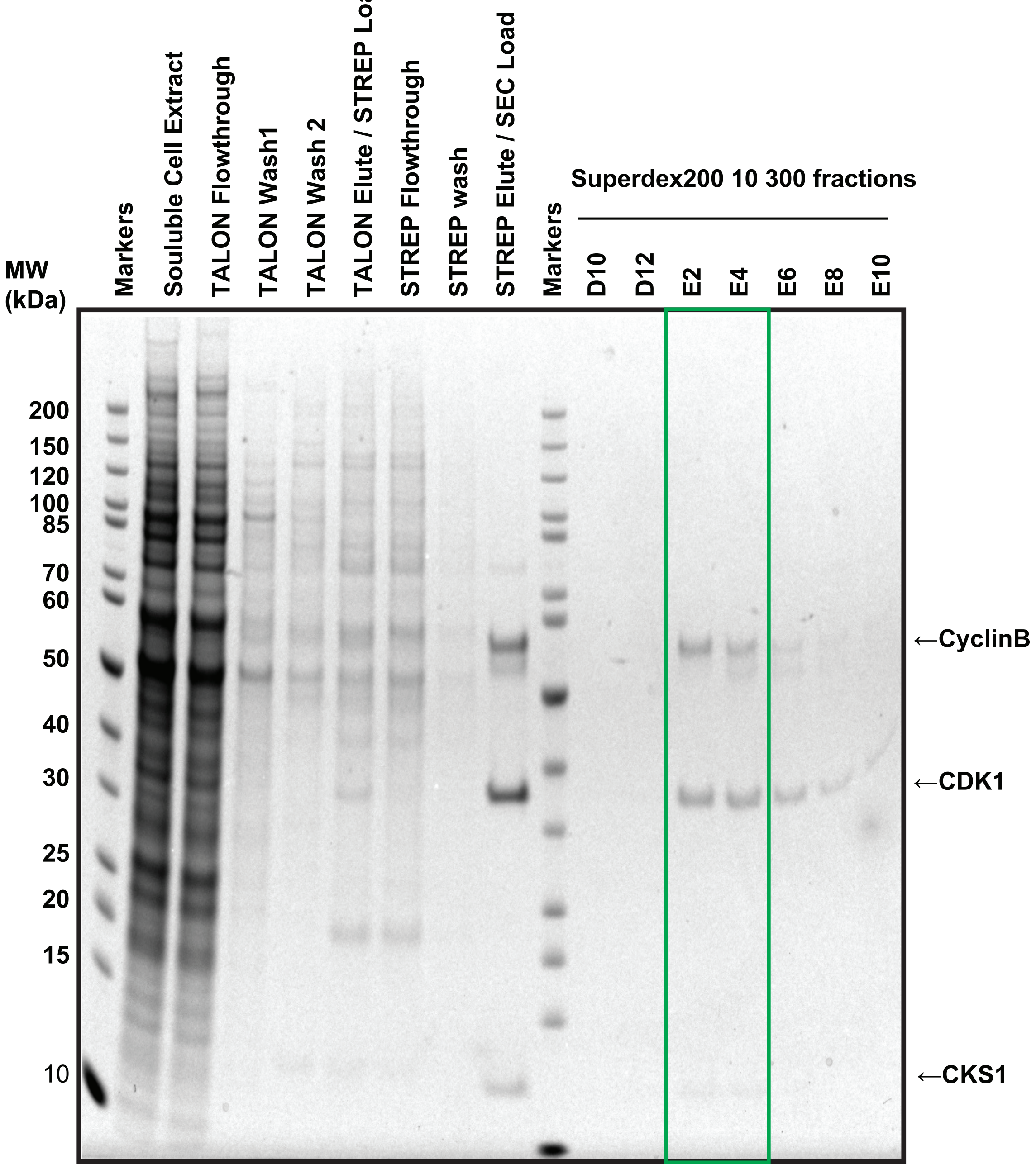
Markers
Souluble Cell Extract
TALON Flowthrough
TALON Wash1
TALON Wash 2
TALON Elute / STREP Load
STREP Flowthrough
STREP wash
STREP Elute / SEC Load
Markers
Superdex200 10 300 fractions
D10
D12
E2
E4
E6
E8
E10
MW (kDa)
200
150
120
100
85
70
60
50
40
30
25
20
15
10
←CyclinB
←CDK1
←CKS1

Figure S4

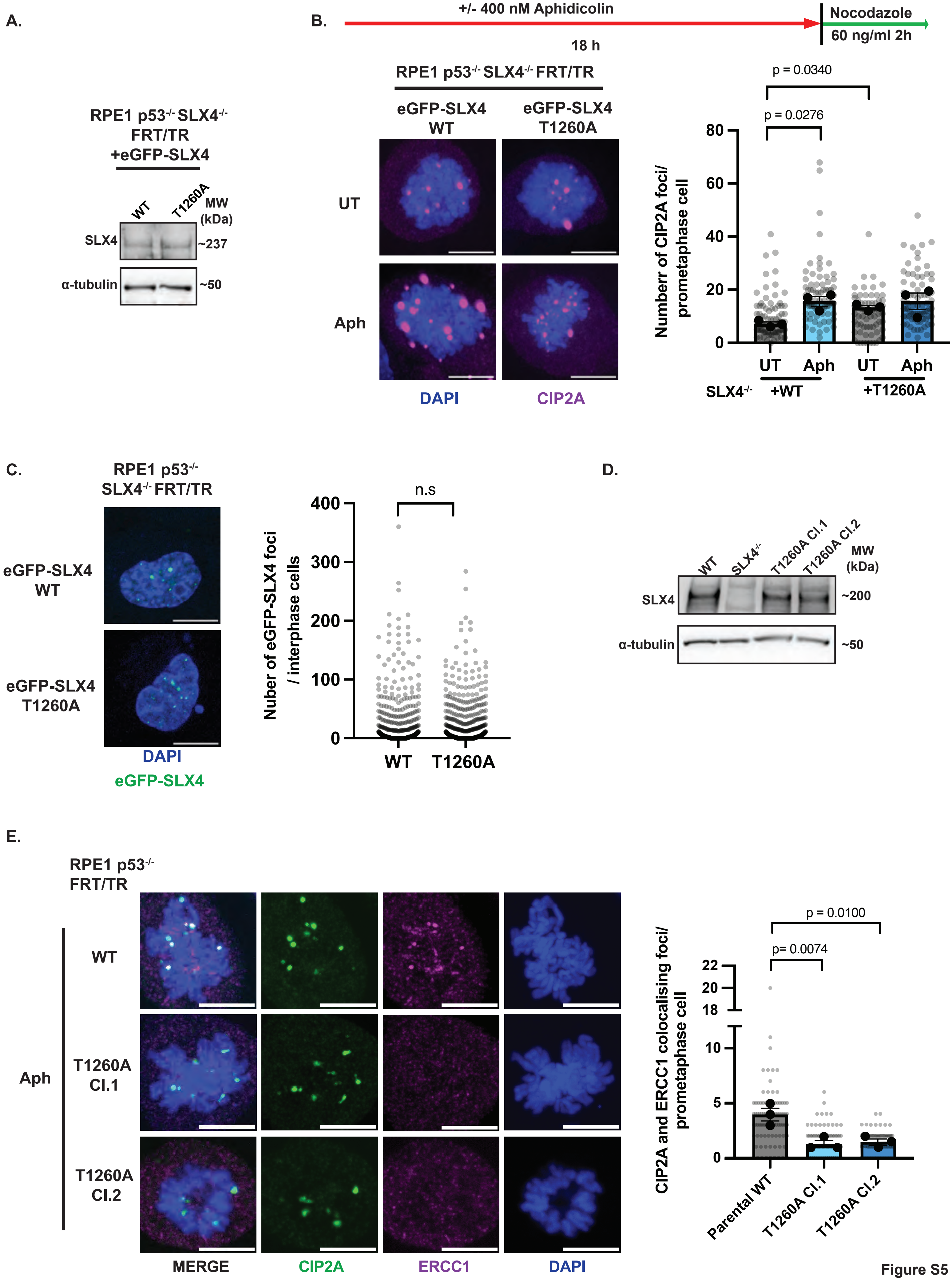

A.
RPE1 p53-/- SLX4-/- FRT/TR +eGFP-SLX4
WT
T1260A
MW (kDa)
SLX4
~237
α-tubulin
~50
B.
+/- 400 nM Aphidicolin
18 h
Nocodazole
60 ng/ml 2h
RPE1 p53-/- SLX4-/- FRT/TR
eGFP-SLX4 WT
eGFP-SLX4 T1260A
UT
Aph
DAPI
CIP2A
p = 0.0340
p = 0.0276
Numberr of CIP2A foci/ prometaphase cell
80
60
40
20
0
UT
Aph
UT
Aph
SLX4-/-
+WT
+T1260A
C.
RPE1 p53-/- SLX4-/- FRT/TR
eGFP-SLX4 WT
eGFP-SLX4 T1260A
DAPI
eGFP-SLX4
n.s
Nuber of eGFP-SLX4 foci / interphase cells
400
300
200
100
0
WT
T1260A
D.
WT
SLX4-/-
T1260A Cl.1
T1260A Cl.2
MW (kDa)
SLX4
~200
α-tubulin
~50
E.
RPE1 p53-/- FRT/TR
Aph
WT
T1260A Cl.1
T1260A Cl.2
MERGE
CIP2A
ERCC1
DAPI
p = 0.0100
p= 0.0074
CIP2A and ERCC1 colocalising foci/ prometaphase cell
22
20
18
10
5
0
Parental WT
T1260A Cl.1
T1260A Cl.2


Figure S5

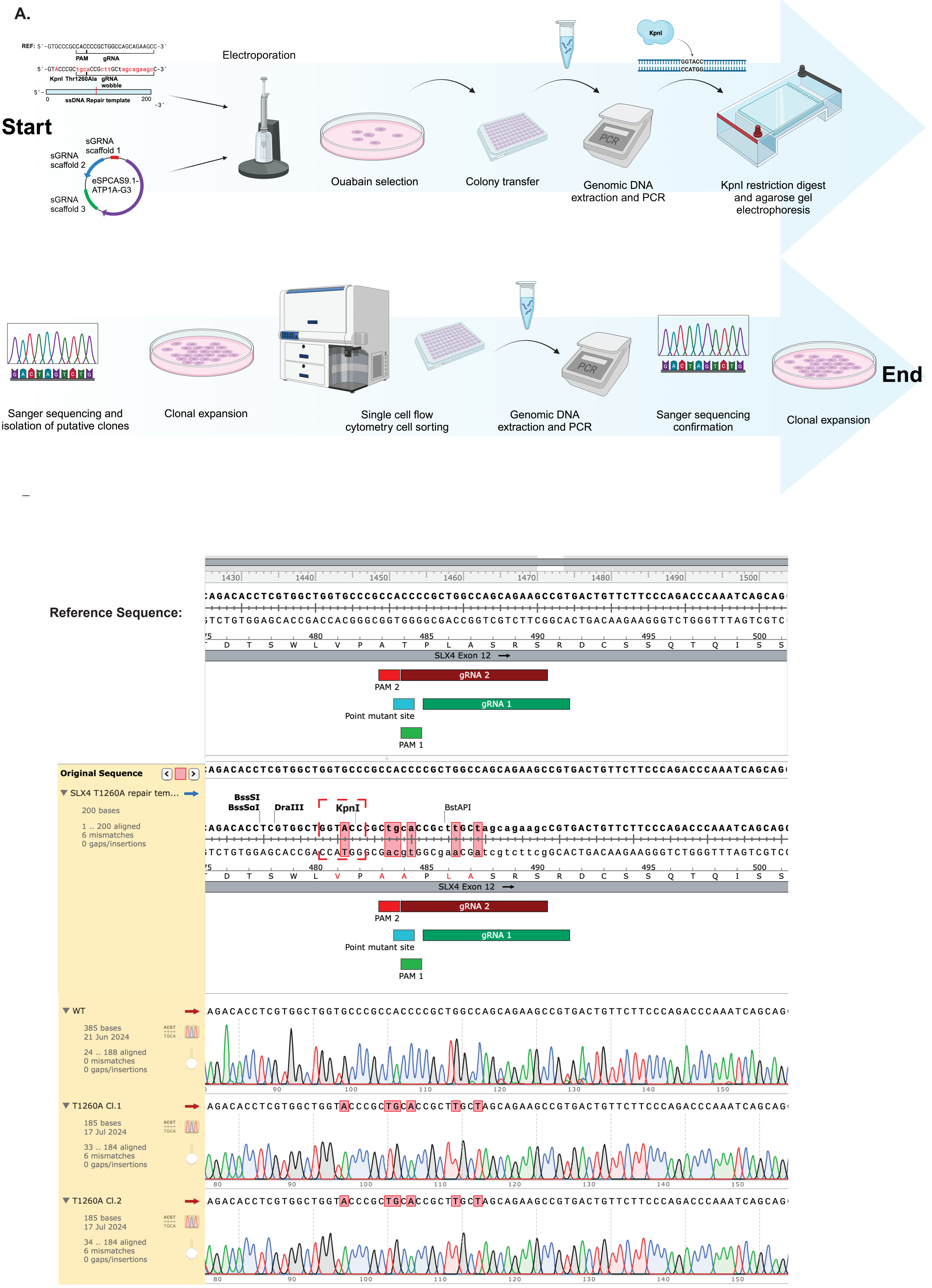
A.
REF: 5'-GTGCCCGCCACCCCGCTGGCCAGCAGAAGCC-3'
PAM
gRNA
Kpnl
Thr1260Ala
gRNA wobble
ssDNA Repair template
Start
sGRNA scaffold 1
sGRNA scaffold 2
sGRNA scaffold 3
eSPCAS9.1-ATP1A-G3
Electroporation
Ouabain selection
Colony transfer
Genomic DNA extraction and PCR
KpnI restriction digest and agarose gel electrophoresis
Sanger sequencing and isolation of putative clones
Clonal expansion
Single cell flow cytometry cell sorting
Genomic DNA extraction and PCR
Sanger sequencing confirmation
Clonal expansion
End
Reference Sequence:
SLX4 Exon 12
gRNA 2
PAM 2
Point mutant site
gRNA 1
PAM 1
Original Sequence
SLX4 T1260A repair tem...
200 bases
1 .. 200 aligned
6 mismatches
0 gaps/insertions
BssSI
BssSaI
DraIII
KpnI
BstAPI
WT
385 bases
21 Jun 2024
24 .. 188 aligned
0 mismatches
0 gaps/insertions
T1260A Cl.1
185 bases
17 Jul 2024
33 .. 184 aligned
6 mismatches
0 gaps/insertions
T1260A Cl.2
185 bases
17 Jul 2024
34 .. 184 aligned
6 mismatches
0 gaps/insertions

Figure S6

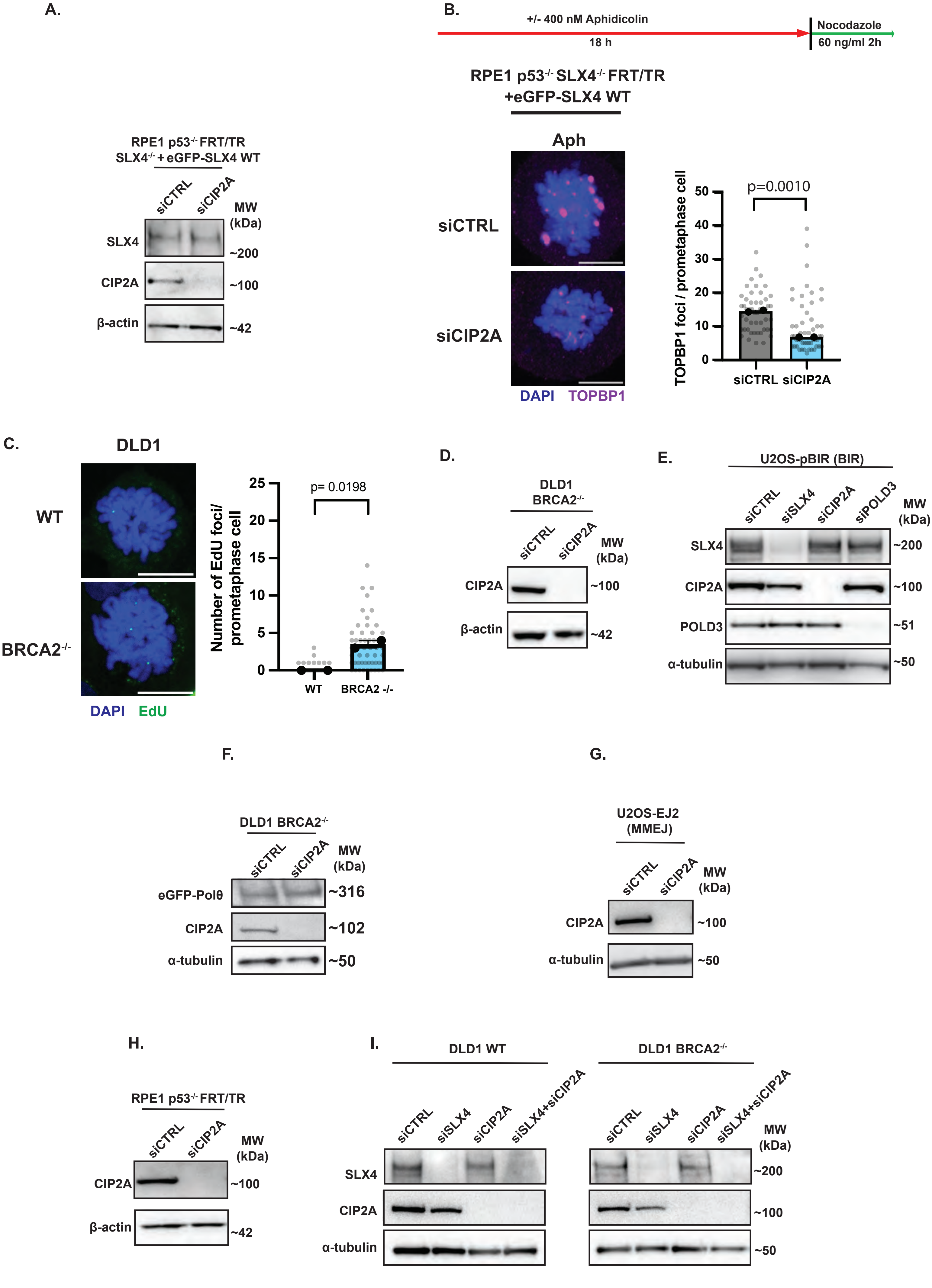
A.
RPE1 p53-/- FRT/TR SLX4-/- + eGFP-SLX4 WT
siCTRL
siCIP2A
MW (kDa)
SLX4
~200
CIP2A
~100
β-actin
~42
B.
+/- 400 nM Aphidicolin
18 h
Nocodazole
60 ng/ml 2h
RPE1 p53-/- SLX4-/- FRT/TR +eGFP-SLX4 WT
Aph
siCTRL
siCIP2A
DAPI
TOPBP1
TOPBP1 foci / prometaphase cell
p=0.0010
50
40
30
20
10
0
siCTRL
siCIP2A
C.
DLD1
WT
BRCA2-/-
DAPI
EdU
Number of EdU foci/ prometaphase cell
p= 0.0198
25
20
15
10
5
0
WT
BRCA2 -/-
D.
DLD1 BRCA2-/-
siCTRL
siCIP2A
MW (kDa)
CIP2A
~100
β-actin
~42
E.
U2OS-pBIR (BIR)
siCTRL
siSLX4
siCIP2A
siPOLD3
MW (kDa)
SLX4
~200
CIP2A
~100
POLD3
~51
α-tubulin
~50
F.
DLD1 BRCA2-/-
siCTRL
siCIP2A
MW (kDa)
eGFP-Polθ
~316
CIP2A
~102
α-tubulin
~50
G.
U2OS-EJ2 (MMEJ)
siCTRL
siCIP2A
MW (kDa)
CIP2A
~100
α-tubulin
~50
H.
RPE1 p53-/- FRT/TR
siCTRL
siCIP2A
MW (kDa)
CIP2A
~100
β-actin
~42
I.
DLD1 WT
DLD1 BRCA2-/-
siCTRL
siSLX4
siCIP2A
siSLX4+siCIP2A
MW (kDa)
SLX4
~200
CIP2A
~100
α-tubulin
~50

Figure S7

A.

B.

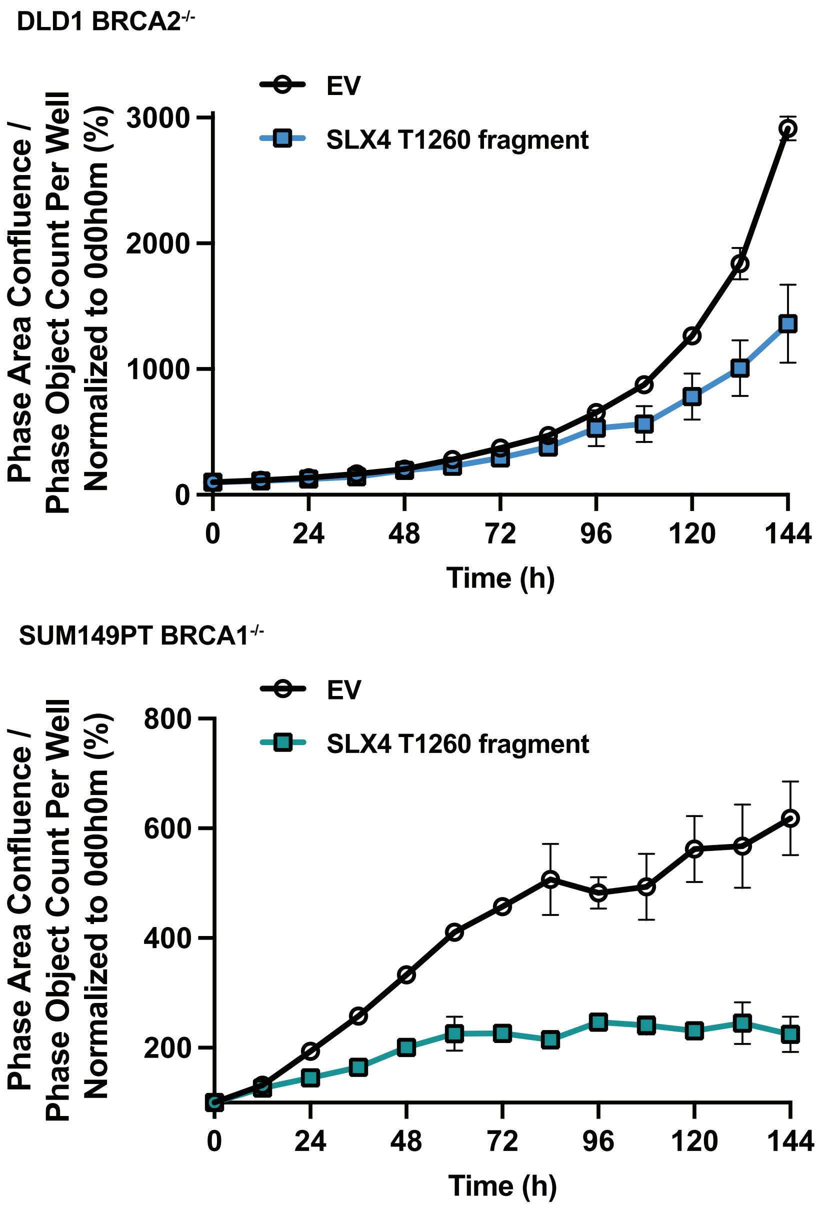
DLD1 BRCA2-/-
EV
SLX4 T1260 fragment
Phase Area Confluence / Phase Object Count Per Well Normalized to 0d0h0m (%)
3000
2000
1000
0
0
24
48
72
96
120
144
Time (h)
SUM149PT BRCA1-/-
EV
SLX4 T1260 fragment
Phase Area Confluence / Phase Object Count Per Well Normalized to 0d0h0m (%)
800
600
400
200
0
24
48
72
96
120
144
Time (h)

Figure S8

### Materials and Methods

#### Cell lines and compound treatments

HCT116-TOPBP1-mAID-Clover cells were a generous gift from Dr D. Cortez. RPE1 $p53^{-/-}$ FRT/TR cells were provided by Dr. S. Jackson. RPE1 $p53^{-/-}$ FRT/TR, HCT116-TOPBP1-mAID-CLOVER and HEK293TN (RRID:CVCL_UL49) cell lines were cultured in Dulbecco's modified Eagle's medium (DMEM) supplemented with 10% fetal bovine serum (FBS) and standard antibiotics. DLD1 (RRID:CVCL_0248), DLD1 $BRCA2^{-/-}$ and SUM149PT (RRID:CVCL_3422) cells were kindly provided by Dr. C. Lord. DLD1 and $BRCA2^{-/-}$ cells were cultured in RPMI-1640 supplemented 10% FBS with the addition of 2 mM L-glutamine and penicillin/streptomycin antibiotics. SUM149PT cells were grown in Ham's F-12 medium supplemented with 5% FBS, 10 µg/mL insulin, 0.5 µg/mL hydrocortisone and penicillin/streptomycin antibiotics. U2OS cells (RRID:CVCL_0042) stably integrated with EJ2 (RRID:Addgene_44025, (*1*)) or (pBIR RRID:Addgene_49807, (*2*)) Dulbecco's modified Eagle's medium (DMEM) supplemented with 10% fetal bovine serum (FBS) and standard antibiotics

Cells were treated with 400 nM Aphidicolin (Sigma Aldrich, A4487), 5 µM ART558 (S9936, Selleckchem), 500 µM 3-indolacetic acid (Sigma Aldrich, I2886) or Nocodazole (Sigma Aldrich, 487928) where indicated. For induction of expression of proteins under tet repressor control cells were treated with 10 ng/ ml (RPE1 $p53^{-/-}$ FRT/TR $SLX4^{-/-}$ system) or 100 ng/ml (DLD1 $BRCA2^{-/-}$) Doxycycline for 24 hours (Sigma Aldrich, D3447) where indicated.

Cell lines were regularly tested to confirm the absence of mycoplasma contamination using the MycoAlert Mycoplasma Detection Kit (Lonza, LT07-318).

#### Plasmids

pcDNA5-Hygro-FRT/TO-GFP-SLX4 was kindly provided by Dr. John Rouse. The SLX4 coding sequence was transferred to pcDNA5-Neo-FRT/TR (RRID:Addgene_41000; a gift from Dr. J. Mansfeld) via HiFi DNA assembly. pcDNA5-Neo-FRT/TO-GFP-SLX4 T1260A plasmid was generated by subjecting the pcDNA5-FRT/TO-Neo-GFP-SLX4 plasmid to site directed mutagenesis using the Q5 site directed mutagenesis kit (NEB). pENTR223.1-POLQ (HsCD00353909**)** was obtained from the DNASU plasmid repository. The POLQ cDNA sequence was amplified from pENTR223.1-POLQ and an eGFP containing fragment was amplified from pcDNA5-FRT/TO-GFP, the fragments were transferred into the pLVX-TET-ONE-Puro (Takara) backbone via HiFi DNA assembly to generate pLVX-TET-ONE-eGFP-POLQ. For the generation of an inducible minimal SLX4 T1260 encompassing fragment, a DNA sequencing encoding an amino acid stretch spanning the surrounding region of SLX4 T1260 was amplified and cloned into pLVX-TET-ONE using HiFi assembly. pOG44 (Thermo Scientific, V6005-20) was used for Flp-Inrecombintation. pVLX-TET-ONE-PURO (Takara), pMD2.G (RRID:Addgene_12259 ) and 4.4µg of psPAX2 (RRID:Addgene_12260) were used for lentiviral work. AIO-mCherry (RRID:Addgene_74120) kindly provided by Dr. S. Jackson, was used as backbone for CRISPR knock-out of SLX4. For amplification, plasmids with the pCDNA5 backbone were transformed and amplified in dH5α bacteria (Thermo Scientific, EC0112) and those used for lentiviral work were transformed and amplified in NEB stable bacteria (NEB, C3040).

#### Inducible cell line generation

To facilitate Flp-In recombination, 2 µg of The Flp-Recombinase expressing plasmid pOG44 was co-transfected with 6 µg pcDNA5-FRT/TO-GFP SLX4 WT/T1260A into RPE1 $p53^{-/-}$

FRT/TR cells, using Lipofectamine 3000 (Invitrogen, L3000001) following manufacturers guidelines. Cells were incubated for 24 hours at 37 °C with 5% $CO_2$ followed by addition of 1 mg/ml G418 (Invivogen, ant-gn) to facilitate selection. G418 was replaced every 72 hours. After 14 days 100 ng/ml of doxycycline was added to the media, and the cells were incubated a further 24 hours at 37 °C with 5% $CO_2$. eGFP expressing cells were sorted by fluorescence activated cell sorting (FACS). Cells were maintained in 500 µg/ ml G418 containing media during continuous culture.

To facilitate lentiviral transduction and integration of inducible eGFP-POLQ, 10µg of pLVX-TET-ONE-eGFP-POLQ was Co-transfected with 1.8µg of pMD2.G ( RRID:Addgene_12259 ) and 4.4µg of psPAX2 (RRID:Addgene_12260) using Lipofectamine 3000 (Invitrogen, L3000001) following manufacturers guidelines into HEK293TN cells. Cells were incubated for 72 hours at 37 °C with 5% $CO_2$ in antibiotic free media. At which point the media was collected, centrifuged at 1500 rpm for 5 min and filtered using a 0.45 µM syringe filter. The filtered lentivirus - containing media was then aliquoted and stored at -80 °C. DLD1 $BRCA2^{-/-}$ cells were subsequently transduced by the addition of lentivirus - containing media and 6 µg/ml Polybrene (Sigma Aldrich, TR-1003). Cells were incubated 24 hours at 37 °C with 5% $CO_2$ in antibiotic free media. Media was then replaced with media containing 4 µg/ml Puromycin (Invivogen, ant-pr) and replaced every 72 hours. After 10 days 100 ng/ ml doxycycline was added to the media and cells were incubated a further 24 hours at 37 °C in 5% $CO_2$ followed by FACS sorting of eGFP positive cells. Cells were maintained in 4 µg/ ml puromycin containing media during continuous culture.

To facilitate lentiviral transduction and integration of inducible a minimal SLX4 T1260 encompassing fragment, 10µg of pLVX-TET-ONE- SLX4 T1260 (fragment) plasmid or pLVX-TET-ONE (Empty vector) was co-transfected with 1.8µg of pMD2.G and 4.4µg of psPAX2 using Lipofectamine 3000 (Invitrogen, L3000001) following manufacturers guidelines into HEK293TN cells. Cells were incubated for 72 hours at 37 °C with 5% $CO_2$ in antibiotic free media. At which point the media was collected, centrifuged at 1500 rpm for 5 min and filtered using a 0.45 µM syringe filter. The filtered lentivirus - containing media was then aliquoted and stored at -80 °C. DLD1 $BRCA2^{-/-}$ cells were subsequently transduced by the addition of lentivirus - containing media and 6 µg/ml Polybrene (Sigma Aldrich, TR-1003). Cells were incubated 24 hours at 37 °C with 5% $CO_2$ in antibiotic free media. Media was then replaced with media containing 4 µg/ml Puromycin (Invivogen, ant-pr) and replaced every 72 hours. After 10 days 100 ng/ ml doxycycline was added to the media and cells were incubated a further 24 hours at 37 °C in 5% $CO_2$ followed by FACS sorting of eGFP positive cells. Cells were maintained in 4 µg/ ml puromycin containing media during continuous culture.

### **Generation of SLX4 knock-out cells by CRISPR/CAS9 nickase**

To facilitate generation of SLX4 knock-out cells previously characterized guide RNA sequence containing complementary oligonucleotides with overhangs compatible with a BbsII or BsaI restriction sites were purchased from IDT.

SLX4gRNA1F Sequence: 5′-ACCG TG TCC CAA AGG ATC CTC AAG-3′
SLX4gRNA1R Sequence: 5′-AAAC CT TGA GGA TCC TTT GGG ACA-3′
SLX4gRNA2F Sequence: 5′-ACCG GT AGG ACC AAT TGT GCT GTG-3′
SLX4gRNA2R Sequence: 5′-AAAC CA CAG CAC AAT TGG TCC TAC CGG T-3′

Subsequently, the oligonucleotides were phosphorylated and annealed by combining 1 µl each of 100 uM complementary oligonucleotides with 1 µl of 10x T4 ligation buffer (NEB, B0202S) with 0.5 µl of T4 PNK (NEB, M0201S) in a total reaction volume of 10 µl. Subsequently, the reaction mixture was incubated at 37 °C for 30 minutes followed by 95°C for 5 minutes and gradient cooled at -5°C/ minute to 25 °C. The annealed and phosphorylated oligonucleotides were then dilute 1:250 in $dH_2O$. Then, 100 ng of AIO-mCherry (RRID:Addgene_74120) kindly provided by Dr. S. Jackson, was digested and ligated by combining 1µl of diluted and annealed phosphorylated oligonucleotide with 1 mM DTT, 1 mM ATP, 0.5 µl of FastDigest BbsI (Thermo Scientific, ER1011), 0.25 µl of T7 DNA ligase (NEB, M0318S) in a final reaction volume of 10 µl. The reaction mixture was then incubated 37 °C for 5 minutes then 23 °C for 5 minutes for 6 cycles. The reaction mixture was then treated with PlasmidSafe exonuclease by combining 5.5 µl of the previous ligation reaction with 1 mM ATP and 0.5 µl of PlasmidSafe exonuclease (LGC Biosearch Technologies, E3101K) in a final reaction volume of 7.5 µl. Subsequently, 3 µl of the treated ligation product was transformed into DH5-Alpha competent cells, plated on LB-Agar/ 50 µg/ml ampicillin containing plates and incubated overnight at 37 °C. Colonies were picked and grown overnight in 3 ml LB-broth containing 50 µg/ml ampicillin. Plasmid was purified using the GeneJET Plasmid Miniprep Kit (Thermo Scientific, K0502). The purified plasmid was then digested and the second pair of annealed and phosphorylated oligonucleotides were ligated as above, but instead using BsaI-HF (NEB, R3733S), followed by PlasmidSafe treatment and transformation into DH5-Alpha cells. Colonies were picked, grown overnight in LB broth as above followed by generation of glycerol stocks and isolation of plasmid DNA. The retrieved plasmid DNA was then sequenced by Source Bioscience using 3.2 pmol of the following sequencing primer:

AIOseq: 5′-CTTGATGTACTGCCAAGTGGGC-3′

Glycerol stocks corresponding to plasmids confirmed to contain both guide RNA sequences were then streaked on LB/Agar plates incubated overnight at 37 °C. Colonies were picked and grow overnight in 50 ml of LB-broth containing 50 µg/ ml ampicillin. Plasmid DNA was then purified using the ZymoPURE II Plasmid Midiprep Kit (Zymo Research, D4201) and resequenced by Source Bioscience using the above AIOseq sequencing primer. Once confirmed RPE1p53$^{-/-}$ FRT/TR cells were transfected with 1.5 µg of plasmid DNA using Lipofectamine 3000 transfection reagent following manufacturer's instructions (Invitrogen, L3000001). After 24 hours incubation at 37 °C with 5% $CO_2$ cells were trypsinised and 10,000 mcherry positive cells were bulk fluorescence activated cell sorted using BD FACSymphony S6 Cell Sorter into a well of a 6 well tissue culture dish, then incubated for a further 7 days at 37 °C with 5% $CO_2$. Subsequently, the single mCherry negative cells were sorted into 96 well tissue culture plates using BD FACSymphony S6 Cell Sorter into 96 well tissue culture plates and incubate at 37 °C with 5% $CO_2$ for 14 days. Colonies were then transferred to wells of a 6 well plates and incubated for a further 5 days at 37 °C with 5% $CO_2$. At this stage, cells were trypsinised and 90% of the cell suspension was isolated for western blot analysis for loss of SLX4 expression at protein level while media was added to the remianing 10 % and incubated at 37 °C with 5% $CO_2$. Once loss of SLX4 protein was detected putative clones were isolated and expanded followed by genomic DNA extraction using the Monarch genomic DNA purification kit (NEB, T3010). Using forward (5′-CAACCACCACCACTACCTAAC-3′) and reverse (5′- GCGAAACCCTGTCTCTACTAAA-3′) primers flanking the edit site and isolated genomic DNA, PCR was carried out using Q5 high fidelity polymerase (NEB, M0491) following manufacturer's instructions. The PCR product was resolved on a 1% agarose/ 1xTBE gel, followed by gel extraction using Zymoclean Gel DNA Recovery Kit (Zymo research, D4002). The purified product was then submitted to Source BioScience for sanger sequencing analysis using the forward PCR primer as a sequencing primer.

**<u>Generation of SLX4 T1260A knock-in cells by CRISPR/CAS9 nuclease marker free co-selection</u>**

RPE1 p53$^{-/-}$ FRT/TR SLX4 T1260A cells, were generated by CRISPR Cas9 mediated knock-in via marker free co-selection, as previously described (*3*). Briefly, two guide RNA sequences were chosen proximal to SLX4 T1260 and assessed for their on-target and off-target efficiency using Benchling and CRISpick guide RNA design tools. Subsequently, a 200 bp repair template was designed to introduce the T1260A point mutation and silent mutations upstream of the PAM sequences to introduce a KPNI restriction endonuclease site. In addition, silent mutations were included within the guide RNA recognition sequences to inhibit further cleavage by the Cas9 sgRNA complex. Previously, Dr. Jörg Mansfeld modified the eSPCAS9.1-ATP1A1-G3 plasmid (RRID:Addgene_86613) a kindly provided by Dr. Yannick Doyon, to include an additional third U6-gRNA cassette with BsmBI sites to create plasmid PLJM787. Complementary single stranded DNA oligos were purchased from IDT to include overhangs compatible with cloning into BbsI and BsmBI sites located proximal to the available encoded gRNA scaffolds.

Guide RNA 1 oligo pair:
5′- CACCGACGGCTTCTGCTGGCCAGCG - 3′
5′- AAACCGCTGGCCAGCAGAAGCCGTC - 3′

Guide RNA 2 oligo pair:
5′- CACCGAGAACAGTCACGGCTTCTGC-3′
5′-AAACGCAGAAGCCGTGACTGTTCTC-3′
T1260A repair template:
5′-
CTCTTTGGGCAGGAGAGGGGCTCCCTGGCTGTTCTGTGACCGTGAGAGCAGCCCCAG
CGAGGCCAGCACCACAGACACCTCGTGGCTGGTACCCGCtgcaCCGcttGCtagcagaagcC
GTGACTGTTCTTCCCAGACCCAAATCAGCAGCCTCAGGAGCGGGCTGGCCGTGCAGG
CGGTGACTCAGCACACGCCCAGGG-3′

ATP1A1 repair template:
5′-
CAATGTTACTGTGGATTGGAGCGATTCTTTGTTTCTTGGCTTATAGCATCAGAGCTGCTA
CAGAAGAGGAACCTCAAAACGATGACGTGAGTTCTGTAATTCAGCATATCGATTTGTAG
TACACATCAGATATCTT-3′

The above RNA encoding oligonucleotides for knock in were phosphorylated and annealed followed by cloning into PLJM787 following the same method described for preparation of AOI-mCherry described in the above method sub-section.

Subsequently, RPE1 p53$^{-/-}$ FRT/TR cells were electroporated using Neon Transfection System. Per electroporation shot 300.000 cells were mixed with 500ng of plasmid, 2pmol of ATP1A1 ALT-R-HDR-Donor oligo (IDT) repair template and 6pmol of the T1260A ALT-R-HDR-Donor oligo (IDT) repair template. Four days after electroporation cells were incubated with media containing 250 nM Ouabain, ATP1A1 inhibitor (Sigma Aldrich, Cat. No: O3125). After 7 days Ouabain resistant clones were picked and transferred into 96-well plates. Each picked clone was duplicated in an additional 96 well plate, and one copy was used for preparation of gDNA. Cells were washed with PBS and lysed in 100µl of: 1mM $CaCl_2$, 3 mM $MgCl_2$, 1 mM EDTA, 10 mM Tris pH 7.4, 1% Triton X-100, supplemented with fresh 0.2 mg/ml Proteinase K (NEB, P8107S). Cells were incubated with lysis buffer at 37°C for 10 minutes and subsequently transferred to a 96 well PCR plate and sealed with plastic plate sealing film.

The plate was transferred to the thermocycler and incubated at 65°C for 10 minutes followed by 95°C 15 minutes. The plate was then stored at -20°C until use.
To PCR amplify the potentially edited region of SLX4 2µl of the gDNA lysate were used for a 25µl standard PCR reaction using Phusion high fidelity polymerase (NEB, M0530S) using the forward (5′-AGCAGGAGGATGAGGGGG-3′) and reverse primers (5′-CCGCCTGCACGGCCA-3′) flanking the edited genomic DNA sequence.  1 µl of KpnI

restriction endonuclease was added to the PCR reaction and incubated for 18 hours at 37 °C. Incorporation of the repair template was verified with restriction digest of the amplified DNA amplicons after agarose gel electrophoresis analysis. Respective clones that demonstrated restriction digestion were subjected to a further round of the above PCR followed by agarose gel electrophoresis and gel purification using the Zymoclean Gel DNA Recovery Kit (Zymo research, D4021). The purified product was then submitted for sanger sequencing analysis by Source Bioscience, using the above forward PCR primer as a sequencing primer, to confirm knock-in.

**Immunoblotting**

Cell lysis was carried out in 1x RIPA buffer (Sigma-Aldrich), supplemented with 1× SIGMAFAST protease inhibitors (Sigma-Aldrich), and 1xPhosStop phosphatase inhibitors (Roche), on ice for 15 min followed by centrifugation at 1.2 × $10^3$ rpm for 20 min followed by collection of supernatant containing cell lysate. Cell lysates were prepared in SDS loading buffer (2% SDS, 10% (v/v) glycerol, 2% 2-Mercaptoethanol and 62.5 mM Tris–HCl, pH 6.8) followed by boiling at 95°C for 10 min. Protein concentrations were determined by the BCA or by the Bradford assay. Samples were resolved by SDS-PAGE and transferred to nitrocellulose membrane followed by blocking in 5% low fat milk in 1× TBS/ 0.1% Tween-20 for 1 h at room temperature. Membranes were washed 3 × 5 min in 1× TBS/ 0.1% Tween-20 and incubated overnight at 4°C in the indicated primary antibodies in 5% low fat milk in 1x TBS/ 0.1% Tween-20. Membranes were subsequently washed 3 × 5 min in 1× TBS/ 0.1% Tween-20 and incubated in 5% low fat milk in 1× TBS/ 0.1% Tween-20 containing secondary antibodies for 1 h at room temperature. Membranes were subsequently washed 3 × 5 min in 1× TBS/ 0.1% Tween-20 and developed using Immobilon Western HRP Substrate (Millipore, WBKLS0S00) and imaged using the Azure C280, 300 or 600 instruments (Azure biosystems).

Primary antibodies used were: α-Tubulin ((Sigma-Aldrich Cat# T5168, RRID:AB_477579, 1:100 000), β-Actin (Sigma-Aldrich Cat# A2066, RRID:AB_476693, 1:1000), GFP (Roche Cat# 11814460001, RRID:AB_390913, 1:500), MUS81 (Santa Cruz Biotechnology, (Santa Cruz Biotechnology Cat# sc-47692, RRID:AB_2147129,1:500), ERCC1 ((Santa Cruz Biotechnology Cat# sc-17809, RRID:AB_2278023,1:500), SLX4 (MRC-PPU Cat# S714C, RRID:AB_2752254, 1:500), SLX4 pT1260 (Genscript). TOPBP1 (Santa Cruz Biotechnology Cat# sc-271043, RRID:AB_10610636,1:500), MDC1 (Abcam Cat# ab11171, RRID:AB_297810, 1:1000), CIP2A (Santa Cruz Biotechnology Cat# sc-80659, RRID:AB_1121640 ,1:500), POLD3 (Abnova Cat# H00010714-M01, RRID:AB_606803, 1:500), TOP3A (Proteintech Cat# 14525-1-AP, RRID:AB_2205881, 1:3000), BLM (Bethyl Cat# A300-110A, RRID:AB_2064794, 1:500).

Secondary antibodies used were anti-mouse IgG-HRP (Dako, P0447, 1:2000), anti-rabbit IgG-HRP (Dako, P0448, 1:5000) and anti-sheep IgG-HRP (Abcam Cat# ab6747, RRID:AB_955453, 1:500).

**Phosphatase incubation of western blot membranes**

After western blot transfer, the membrane was washed 3 x for 5 minutes n 1x TBS/ 0.1% Tween-20 (TBS-T). The western blot membrane was divided in half by vertical cutting. The divided membrane was then separated into two 50 ml centrifuge tubes containing 10 ml phosphatase reaction buffer (NEB, 1x rCutSmart buffer B6004SVIAL) supplemented with 1 mM $MgCl_2$. To one tube 1 µg/ml of lambda protein phosphatase and 1 unit/ml of shrimp alkaline phosphatase (rSAP; NEB, M0371). The tubes were then placed in a rotating hybridization oven at 30°C for 1 hour followed by 2 hours at 37 °C. The membranes were then washed once in TBS-T, followed by blocking with 5% low fat milk/ TBS-T for 1 hour.

Membranes were then incubated in with antibodies and developed as described in the above immunoblotting methods.

**RNA interference.**

To facilitate transient depletion of TOPBP1, SLX4 or CIP2A, cells were transfected with 30 pmol of oligonucleotides using Lipofectamine RNAimax transfection reagent (Invitrogen, 13778100), according to the manufacturer's reverse transfection protocol. siRNA targeting luciferace was used as a non-targetting control siRNA. For experiments involving CIP2A ON-TARGETplus siRNA SMARTpool containing a mixture of four oligonucleotides was used ( Horizon, L-014135-01-0005). For POLD3 RNAi a SMARTpool of four oligonucleotides was used ( Horizon, L-026692-01-0005). For depletion of Polθ by RNAi a SMARTpool consisting of four oligonucleotides was used ( Horizon, L-015180-01-0005) For experiments involving SLX4 siRNA, SLX4 (1) and (2) oligonucleotides were mixed at equimolar ratios in advance of use. First pulse of siRNA was followed with second pulse after 24 h following manufacturer’s forward transfection protocol and all subsequently experiments were performed from 72 h post first siRNA pulse at maximum knock down-efficiency.

Small interfering RNA used in this study were as follows:

siRNA targeting luciferase (siCTRL/siLUC): 5′-CGTACGCGGAATACTTCGA-3′

KIAA1524 (CIP2A) ID: L-014135-01-0005 (Horizon) consisting of:

CIP2A (1): 5′-ACAGAAACUCACACGACUA-3′

CIP2A (2): 5′-GUCUAGGAUUAUUGGCAAA-3′

CIP2A (3): 5′-GAACAAGGUUGCAGAUUC-3′

CIP2A (4): 5′-GCAGAGUGAUAUUGAGCAU-3′

SLX4 (1): 5′-GCACAAGGGCCCAGAACAA-dT-dT-3′

SLX4 (2): 5′-GCACCAGGUUCAUAUGUA-dT-dT-3′

TOPBP1: 5′-GUAAAUAUCUGAAGCUGUAUU-3′

POLD3 L-026692-01-0005

Polθ: L-015180-01-0005

**Immunofluorescence microscopy**

For detection of EdU incorporation in mitosis and the analysis of recruitment of TOPBP1, CIP2A, MUS81, ERCC1, eGFP tagged SLX4 and eGFP tagged Polθ, cells were grown on coverslips and treated as described. Subsequently cells were simultaneously fixed and permeabilised in PTEMF buffer (20 mM PIPES pH 6.8, 10 mM EGTA, 0.2% Triton X-100, 1 mM MgCl2 and 4% formaldehyde) for 10 minutes at room temperature. For detection of micronuclei, cells were fixed in 250 mM HEPES pH 7.5, 0.1% Triton X-100, 4% PFA in PBS for 20 minutes at 4 °C.

Coverslips were then incubated for 5 minutes in 1x PBS with three buffer changes followed by incubation with 0.5% Triton X-100/ PBS for 10 minutes at room temperature. Coverslips were then incubated for 5 minutes in 1x PBS with three buffer changes. For EdU detection coverslips were transferred to a humidified chamber and incubated in EdU click it reaction buffer containing alexafluor 488 azide for 1 hour at room temperature followed by three 5 min washes in 1x PBS. Coverslips were then incubated in 5% FBS/PBS for 1 hour followed by overnight incubation at 4 °C with primary antibody diluted in 5% FBS/PBS. Coverslips were then washed 3x 5 minutes in 1x PBS followed by 1 hour incubation at room temperature in 5% FBS/PBS containing secondary antibodies. Coverslips were incubated a further three times for 5 minutes in 1x PBS. Coverslips were then mounted on superfrost microscopy slides using Vectashield mounting medium with DAPI (Vector laboratories, H1200).

Primary antibodies used for immunofluorescence microscopy were: mouse anti-MUS81 (Santa Cruz Biotechnology Cat# sc-53382, RRID:AB_2147138,1:500), mouse anti-ERCC1 (Santa Cruz Biotechnology Cat# sc-17809, RRID:AB_2278023, ,1:500), mouse anti-TOPBP1 (Santa Cruz Biotechnology Cat# sc-271043, RRID:AB_10610636, 1:500), rabbit anti-TOPBP1 (Bethyl Cat# A300-111A, RRID:AB_2272050,,1:500) mouse anti-CIP2A (Santa Cruz Biotechnology Cat# sc-80659, RRID:AB_1121640, 1:500), rabbit anti-CIP2A (Proteintech Cat# 23199-1-AP, RRID:AB_2918079,1:500), or GFP-Booster Alexa Fluor® 488 (ChromoTek Cat# gb2AF488-10, RRID:AB_2827573, 1:500).

Secondary antibodies used for immunofluorescence microscopy were: Alexa Fluor® 647 AffiniPure™ Donkey Anti-Mouse IgG (H+L) (Jackson ImmunoResearch Labs Cat# 715-605-150, RRID:AB_234086, 1:400), Goat anti-Mouse IgG (H+L) Cross-Adsorbed Secondary Antibody, Alexa Fluor™ 647 (Molecular Probes Cat# A-21235, RRID:AB_2535804,1:400), Donkey anti-Rabbit IgG (H+L) Highly Cross-Adsorbed Secondary Antibody, Alexa Fluor™ 647 (Molecular Probes Cat# A-31573, RRID:AB_2536183,1:400) or Donkey anti-Rabbit IgG (H+L) Highly Cross-Adsorbed Secondary Antibody, Alexa Fluor™ 488 (Thermo Fisher Scientific Cat# A-21206, RRID:AB_2535792, 1:400).

Images were acquired using a Zeiss Axio Observer Z1 Marianas™ Microscope attached with a CSU-W spinning disk unit using either a Hamamatsu Flash 4 CMOS camera or a Photometrics Prime 95b sCMOS camera built by Intelligent Imaging Innovations (3i) or a Zeiss Axio Observer Z1 Marianas™ Microscope attached with a CSUX1 spinning disk unit and Hamamatsu Flash 4 CMOS camera built by Intelligent Imaging Innovations (3i). Quantification was carried out using FIJI (ImageJ, RRID:SCR_003070)
) software, CellProfiler (Broad Institute) or using Aivia (Leica Microsystems). Images were analysed in 3D with maximum Z projections used for representative images in the manuscript to assist with visualization.

### EdU labelling

RPE1 p53$^{-/-}$ FRT/TR cells were grown of coverslips and exposed to 400 nM aphidicolin to induce mild DNA replication stress for 18 hours at 37 °C in 5% $CO_2$. Aphidicolin was then washed out by two media replacements followed by a third media replacement containing 20 µM EdU and 60 ng/ml nocodazole and incubated at 37 °C in 5% $CO_2$ for a further 30 minutes. DLD1 WT and DLD1 BRCA2$^{-/-}$ cells were grown on coverslips for 24 hours at 37 °C in 5% $CO_2$ followed by incubation with media containing 20µM EdU for 30 minutes. Cells were then fixed in PTEMF buffer at room temperature for 10 minutes, followed by three 5 minute PBS washes. Subsequently, coverslips were incubated for 10 minutes in 0.5% Triton-X/PBS at room temperature, followed by three 5 minute PBS washes. Coverslips were then mounted using Vectashield mounting medium containing DAPI (Vector laboratories, H1200).

Images were acquired using a Zeiss Axio Observer Z1 Marianas™ Microscope attached with a CSU-W spinning disk unit using either a Hamamatsu Flash 4 CMOS camera or a

Photometrics Prime 95b sCMOS camera built by Intelligent Imaging Innovations (3i). Quantification was carried out using FIJI (ImageJ, RRID:SCR_003070) software.

#### Cellular proliferation analysis by live cell imaging

To analyse cellular proliferation of DLD1 BRCA2$^{-/-}$ cells were treated with siRNA as indicated. At 72 hours post-transfection of the first pulse of siRNA, 1500 cells were seeded per well in a PhenoPlate 96-well optically clear tissue culture plate (Revvity, 6055302). Cells were evenly distributed within the well by pipetting and then incubated at 37 °C with 5 % $CO_2$ for 24 hours. The following day the media was aspirated and replaced with 200 µl of RPMI 1640 media containing standard antibiotics, 10% FBS and 2 mM L-glutatmine, plates were then transferred to an Incucyte S3 Live Cell Analysis System (Sartorius) imaging system and phase images were captured every 12 hours for a total of 120 hours. Subsequently, proliferation was analysed using the Incucyte analysis software, normalising to confluence (%) at the assay start point. Data was exported to Microsoft excel and values were plotted in GraphPad Prism 10.

To analyse cellular proliferation of RPE1 p53$^{-/-}$ FRT/TR and RPE1 p53$^{-/-}$ FRT/TR SLX4 T1260A Cl.1 and 2 cells, 500 cells per well of a 96 well µclear plate (Griener Bio-One, 655096) were seeded in a final volume of 100 µl of DMEM media containing standard antibiotics and 10% FBS and incubated 6 hours at 37 °C with 5 % $CO_2$. Then 2x working dilutions of DMSO, ART558, aphidicolin + DMSO and aphidicolin + ART558 were prepared, then 100 µl of drug dilution was added to each well of the plate to achieve final working concentrations of 5 µM ART558 and 400 nM aphidicolin. Cells were then transferred to an Incucyte SX5 Live Cell Imaging and Analysis System (Sartorius) imaging system and phase images were captured every 12 hours for a total of 120 hours. Subsequently, proliferation was analysed using the Incucyte analysis software, normalising to confluence (%) at the assay start point. Data was exported to Microsoft excel and values were plotted in GraphPad Prism 10.

To analyse cellular proliferation of DLD1 BRCA2$^{-/-}$ or SUM149PT cells, 1500 cells were seeded per well in a PhenoPlate 96-well optically clear tissue culture plate (Revvity, 6055302). Cells were evenly distributed within the well by pipetting and then incubated at 37 °C with 5 % $CO_2$ for 24 hours. The following day the media was aspirated and replaced with 200 µl of RPMI 1640 media containing standard antibiotics, 10% FBS and 2 mM L-glutatmine (DLD1 BRCA2$^{-/-}$ SLX4 T1260 fragment cells or empty vector control expressing) or S Ham's F-12 medium supplemented with 5% FBS, 10 µg/mL insulin, 0.5 µg/mL hydrocortisone and and penicillin/streptomycin antibiotics for SUM149PT (SLX4 T1260 fragment or empty vector control expressing)., plates were then transferred to an Incucyte S3 Live Cell Analysis System (Sartorius) imaging system and phase images were captured every 12 hours for a total of 144 hours. Subsequently, proliferation was analysed using the Incucyte analysis software, normalising to confluence (%) at the assay start point divided by the by the number of objects detected. Data was exported to Microsoft excel and values were plotted in GraphPad Prism 10.

#### Collection of asynchronous, S-phase and M-phase synchronised cells

To facilitate enrichment of HEK293TN cells in S phase or M phase, $2x10^6$ cells were seeded in 15cm tissue culture dishes and incubated at 37 °C with 5% $CO_2$ for 24 hours. For enrichment of S-phase cells, cells were subsequently incubated with media containing 2 mM thymidine (Sigma Aldrich, T9250) for 18 hours. Media was replaced by 3 buffer changes and then cells were incubated for 9 hours at 37 °C with 5% $CO_2$ to facilitate cell cycle progression. The media was then replaced with 2 mM thymidine containing media and cells were incubated for a further 15 hours at 37 °C with 5% $CO_2$. Thymidine containing media was then removed followed by two further media changes to facilitate thymidine wash-out. Cells were incubated for a further 3 hours at 37 °C with 5% $CO_2$ and harvested by trypsinisation followed by inactivation by addition of serum containing media then by centrifugation at 300 rcf for 3 minutes. The supernatant was aspirated, and cells were snap frozen on dry ice. For enrichment of mitotic cells, after 24 hours incubation at 37 °C with 5% $CO_2$ media was replaced with media containing 100 ng/ml nocodazole and incubated at 37 °C with 5% $CO_2$ for 18 hours. At which point the media was collected, followed by 1x PBS wash, collection and then trypsinisation followed by inactivation by addition of serum containing media. Cells were pelleted by centrifugation at 300 rcf for 3 minutes, the supernatant was aspirated, and cells were snap frozen on dry ice. Asynchronous cells were collected in parallel by trypsinisation followed inactivation by addition of serum containing media, then centrifugation at 300 rcf for 3 minutes. Then the supernatant was aspirated, and cells were snap frozen on dry ice. In all cases before centrifugation a 500 µl aliquot of suspended cells was fixed in ice cold 70% ETOH followed by propidium iodide staining and flow cytometry analysis to confirm cell cycle stage by DNA content analysis (see below).

**Flow Cytometry**

Cells resuspended in media after trypsinisation were centrifuged at 300 rcf for 3 minutes. The media was aspirated, and the cell pellet was resuspended in 1ml of 1x PBS. The cells were again pelleted by centrifugation at 300 rcf for 3 mins and the PBS was removed. The cell pellet was then loosened by flicking the tube and 70% ice cold ethanol was added slowly while continually vortexing the sample at ~1000 RPM. The sample was then placed on ice for 30 minutes and stored until the day of processing at -20 °C. On the day of analysis, the cells were pelleted by centrifugation at 300 rcf for 3 mins. The 70% ethanol was aspirated and 1 ml of 1x PBS was added. The sample was centrifuged again at 300 rcf for 3 mins and the PBS was aspirated. The previous PBS wash step was carried out twice more. The sample was then resuspended in 2 ml of 1x PBS with 1 µg/ml RNAse A (Roche, RNASEA-RO- 10109142001) and 1 µg/ml of propidium iodide. The sample was incubated in the dark at 4°C for 4 hours, then vortexed and transferred to a 5 ml flow cytometry tube through a cell strainer. The sample was then analysed using the BD LSR II Analyzer flow cytometry instrument. Cells were gated by analysis of forward area and side scatter area, followed by gating of single cells by gating forward scatter height against forward scatter area. A histogram was then plotted of the PE: TxRed-H PI intensity per event. The data was then analysed using FlowJo (FlowJo LCC) analysis software comparing 7915 events per condition.

**Co-immunoprecipitation**

For TOPBP1 Co-immunoprecipitation experiments, 50µl of Dynabead protein G slurry were washed 3x in (IP2 200 mM NaCl, 0.2% Igepal CA-630, 1 mM MgCl2, 10% glycerol, 5 mM NaF, 2 mM EDTA, 50 mM Tris-HCl, pH 7.5) Beads were then blocked in IP2 + 1% BSA + 1x SIGMAFAST protease inhibitors (Sigma-Aldrich) for 1 h at RT. Blocking buffer was removed and replaced with diluted 1ml IP2 buffer with either 3 µg of anti-TOPBP1 (A300-111A, Bethyl) or 3 µg of anti-rabbit IgG control (AP132, Sigma-Aldrich). The beads and antibody mixture were rotated at 4 °C overnight. Cell pellets were removed from -80 storage and thawed on ice in 2.4 ml IP buffer 1 (100 mM NaCl, 0.2% Igepal CA-630, 1 mM MgCl2, 10% glycerol, 5 mM NaF, 50 mM Tris-HCl, pH 7.5), supplemented with complete PhosSTOP phosphatase inhibitor (PHOSS-RO, Roche), 1x SIGMAFAST protease inhibitor and 25 U ml−1 Benzonase (Novagen). Cells pellets were resuspended, rotated and lysed at 4°C on carousel for 90 min. Benzonase was then inhibited by adjusting the NaCl concentration to 200 mM (48 ml from 5M NaCl stock) and 2 mM EDTA (9.6 ml from 500 mM EDTA stock) and rotated for a further 30 min at 4°C on a carousel. Cell suspensions were then centrifuged at 16,000 rcf for 25 min at 4°C. Soluble supernatant (lysate) from the above centrifugation then transferred to individual 15 ml conical tube. BCA assay was then carried out on to determine protein concentration. Antibody mix is removed from Dynabeads and washed 3x in 500 µl IP2. The antibody bound Dynabeads were transferred to 2ml tubes and IP2 buffer was added to bring each sample to 1 mg/ ml final. The tubes were then rotated at 4°C on carousel for 120 min. The supernatant was then removed while the beads and 500 ml IP2 buffer was added to wash the beads. This step was repeated once. IP2 buffer was then removed, and beads were washed with 500 µl PBS. The PBS was then removed, and the PBS was step was repeated once more. The PBS was removed and discarded and beads bound by antibody-protein complexes were stored at –20 °C prior to IP-MS analysis.

**Mass-spectrometry proteomics**

Beads were re-suspended in 100 µL of 100 mM TEAB, reduced with 10 µL of 50 mM TCEP, and alkylated with 5 µL of 200 mM freshly prepared iodoacetamide. The mixture was incubated at room temperature for 30 minutes in the dark. Proteins were digested overnight with trypsin (500 ng/µL in 0.1% formic acid) at 37°C with shaking. The digested peptides were collected, dried, and cleaned using the Pierce™ High pH Reversed-Phase Peptide Fractionation Kit (Thermo Scientific). TMT-labeled samples were combined, vacuum-dried, fractionated using the same kit, and dried again before MS analysis. Samples were re-suspended in 0.1% TFA prior to analysis.
For label-free samples, LC-MS analysis was performed using a Dionex UltiMate 3000 UHPLC system coupled to an LTQ Orbitrap Lumos mass spectrometer (Thermo Scientific). Chromatographic separation was achieved on an EASY-Spray C18 column (75 µm × 50 cm, 2 µm) at 50°C. The mobile phase consisted of 0.1% formic acid (A) and 80% acetonitrile with 0.1% formic acid (B). The gradient elution was: 0–150 minutes up to 38% B, 150–160 minutes up to 95% B, 160–165 minutes isocratic at 95% B, 165–175 minutes re-equilibration to 5% B, and 175–185 minutes isocratic at 5% B.

Data-independent acquisition (DIA) was performed on a high-resolution Orbitrap mass spectrometer in positive ion mode. An MS1 survey scan was acquired between 8 and 109 minutes at 60,000 resolution with a scan range of m/z 400–900. Quadrupole isolation was used with a Standard AGC target and a custom maximum injection time of 300 ms. DIA was conducted using 42 variable-width isolation windows across m/z 400–900, with precursor ions fragmented by HCD at 30% collision energy. Fragment ions were detected in the Orbitrap at 15,000 resolution over m/z 145–1450. DIA windows were calculated to ensure complete coverage and minimal overlap of the precursor mass range.

For DIA-NN searches, raw data files were deconvoluted and converted to mzML files using MS Converter with filter settings peakPicking (vendor msLevel=1-) and titleMakerData. The data was searched against human proteome (UniProtKB). The analysis employed a library-free workflow. For precursor ion generation, the options FASTA digest for library-free search/library generation and Deep learning spectra, RTs, and IMs prediction were enabled. The digestion parameters allowed for a maximum of one missed cleavage by trypsin, with a maximum of two variable modifications per peptide. Carbamidomethylation of cysteine residues was specified as a fixed modification, while methionine oxidation. Peptide length was restricted to a range of 7 to 30 amino acids. The precursor charge range was set from 2 to 4, with the m/z range for both precursors and fragments set from 300 to 1300. The mass accuracy for both parent and fragment ions was manually adjusted to 10 ppm. A false discovery rate (FDR) threshold of 1% was applied at both the protein and peptide levels, and the match between runs feature was enabled to improve peptide identification across different samples.

TMT-labelled samples were analysed using a Thermo Orbitrap Ascend mass spectrometer. MS1 scans were performed over a mass range of m/z 400–1600 at 120,000 resolution in the

Orbitrap, with standard AGC settings and automatic injection times. Ions with charge states +2 to +6 were included. Dynamic exclusion was set to 45 seconds with a repeat count of 1, a ±10 ppm mass tolerance, and isotopes were excluded from further analysis.
MS2 spectra were acquired in the ion trap using a Turbo scan rate, with 32% HCD collision energy and a maximum injection time of 35 ms. Real-time database searching against Homo sapiens (canonical and isoforms) was conducted using the Comet search engine, considering tryptic peptides with a maximum of 1 missed cleavage. Static modifications were set for carbamidomethylation of C (+57.0215 Da) and TMTpro labelling on K and N-termini (+304.207 Da). Variable modifications included deamidation of N/Q (+0.984 Da) and oxidation of M (+15.9949 Da), allowing up to 2 variable modifications per peptide. A close-out feature was enabled, limiting to 4 peptides per protein.

SPS10-MS3 scans were performed on selected precursors using the Orbitrap at 45,000 resolution with 55% HCD collision energy, a 200% normalized AGC target, and a 200 ms maximum injection time. Data were collected in centroid mode with single micro-scan acquisition. For protein identification and quantification, Proteome Discoverer 3.0 (Thermo Scientific, Proteome Discoverer, RRID:SCR_014477) with SequestHT and Comet search engines was used. Spectra were searched against the UniProt (RRID:SCR_002380) Homo sapiens database with a precursor mass tolerance of 20 ppm and a fragment mass tolerance of 0.02 Da. Peptides were considered fully tryptic, allowing up to 2 missed cleavages. Static modifications included TMT at N-termini/K and carbamidomethylation at C residues. Dynamic modifications included methionine oxidation and deamidation of N/Q. Peptide confidence was assessed using Percolator with a 1% FDR using target-decoy validation. Quantification employed the TMT quantifier node with a 15 ppm integration window using the most confident centroid peak at the MS2 level. Only unique peptides with a signal-to-noise ratio >3 were considered for quantification. Data were normalized to total protein loading, and relative abundances were calculated by dividing normalized values by the average abundance across all TMT channels per biological replicate.
Statistical analysis and data visualisation was performed using Python (RRID:SCR_008394). Gene ontology enrichment analysis was performed using ShinyGO (PMID: 31882993).

**GFP-Trap® Agarose Co-immunoprecipitation**

HEK293TN cells were seeded to be ~25-30% confluence the next day in a 15 cm tissue culture dish. The following day cells were transfected using Lipofectamine 2000 transfection reagent (ThermoScientific) with 24 µg of plasmid DNA encoding eGFP-SLX4 or eGFP-TOPBP1 expression casettes following manufacturer's instructions.
48 hrs post transfection cells were harvested after treatments as described and snap frozen on dry ice and strored at -80 °C. Cell pellets were removed from -80 storage and thawed on ice in 2.4 ml IP buffer 1 (100 mM NaCl, 0.2% Igepal CA-630, 1 mM MgCl2, 10% glycerol, 5 mM NaF, 50 mM Tris-HCl, pH 7.5), supplemented with EDTA Free SIGMAFAST protease inhibitor (Sigma Aldrich) and 25 U $ml^{-1}$ Benzonase (Novagen). Cells were resuspended and rotated at 4°C for 90 min. Benzonase was then inhibited by adjusted NaCl concentration to 200 mM and 2 mM EDTA and rotated for a further 30 min at 4°C on a carousel. Cell suspension was then centrifuged at 16,000 rcf for 25 min at 4°C. Then, 20 µl binding control bead slurry was washed with 500 µl IP buffer 2 (200 mM NaCl, 0.2% Igepal CA-630, 1 mM MgCl2, 10% glycerol, 5 mM NaF, 2 mM EDTA, 50 mM Tris-HCl, pH 7.5). The binding control beads were then centrifuged at 2500 rcf for 2 min. The supernatant was then removed and the IP2 wash was repeated twice more. Cell lysates were then added to the agarose binding control beads in 15 ml centrifuge tubes and subsequently rotated at 4°C on carousel for 60 min. The agarose binding control beads were then pelleted by centrifugation at 2000 rcf for 2 minutes, the supernatant was then transferred to a new tube. BCA assay was then carried out to determine the protein concentration of the pre-cleared cell lysate. Subsequently, 20 µl GFP_TRAP_A bead slurry was washed with 500 µl IP buffer 2 (200 mM NaCl, 0.2% Igepal

CA-630, 1 mM MgCl2, 10% glycerol, 5 mM NaF, 2 mM EDTA, 50 mM Tris-HCl, pH 7.5). The beads were then centrifuged at 2500 rcf for 2 min. The supernatant was removed and the IP2 wash step is repeated twice more. Cell lysates were diluted to 1 mg/ml final concentration in IP buffer 2 and added to the washed GFP_TRAP_A beads in 15 ml conical tubes and rotated at 4°C on carousel for 120 min. The 15 ml conical tubes were then centrifuged at 2000 rcf for 2 min at 4°C and the supernatant was removed. The beads were then washed with 500 µl IP2 buffer and transferred to a 1.5 ml microcentrifuge tube, then centrifuged at 2500 rcf for 2 min at 4°C. The supernatant was removed and and the IP2 wash was repeated twice more. Bound protein complexes were eluted from the the GFP_TRAP_A beads by addition of 50 µl 2x SDS buffer ( 120 mM Tris/Cl pH 6.8, 20 % glycerol, 4 % SDS, 0.04 % bromophenol blue, 10 % β-mercaptoethanol) followed by incubation at 95 °C for 10 min. Tubes were then placed on ice for 10 min followed by centrifugation for 2500 rcf for 2 min and the supernatant was transferred to a new 1.5 ml microcentrifuge tube ready for downstream western blot analysis.

**BRCT 1 recognition motif bioinformatics search**

Consensus motifs for BRCT1 and 2 of TOPBP1 (*4*) were scanned over the SLX4 sequence using the ExPASy ScanProsite server (*5*). Potential sties were checked against Phosphosite plus (*6*) to see if they were documented phosphophorylation sites and the sequence conservation of the sites was also checked using the Proviz Server(*7*).

**Alphafold3 modelling**

Amino acid sequences for TOPBP1(1-300) and SLX4(1240-1280) with pT1260 modification were submitted to the Alphafold3 web server (*8*). Images of the complex, including superposition with known TOPBP1-RAD9 structure, were produced using PyMOL (version2.2.2 ; PyMOL, RRID:SCR_000305)

**TOPBP1(1-290) purification**

BL21(DE3) cells were transformed with pGEX-6P-1 TOPBP1(1-290) WT and K155E, K250E and K155E/K250E mutant variants for protein expression. Cell pellets were resuspended in lysis buffer containing 25 mM HEPES pH 7.5, 200 mM NaCl and 0.5 mM TCEP supplemented with 50U Turbo DNase, disrupted by sonication, and the resulting lysate clarified by centrifugation at 36,000 x *g* for 60 min at 4°C. The supernatant was applied to a 5 ml HiTrap GST column, then washed with buffer containing 50 mM HEPES pH 7.5, 1000 mM NaCl, 0.25 mM TCEP, before retained protein was eluted by application of the lysis buffer supplemented with 20 mM glutathione. This was concentrated using a Vivaspin with a 30 kDa MWCO and applied to a Superdex 200 16/60 size exclusion column equilibrated in 25 mM HEPES pH 7.5, 150 mM NaCl, 1 mM EDTA, 0.5 mM TCEP, 0.002% (v/v) Tween-20.

**CDK1-CyclinB-CKS1 purification**

A single construct containing CDK1, CyclinB and Cks1 was produced using the biGBac system (*9*). SF9 cells were transfected with this bacmid to produce baculovirus for protein expression, and following two rounds of viral amplification, cells were infected at a cell density of 1.5x10^6 using 5% viral stock for expression and harvested after 48 hours. Cell pellets were resuspended in lysis buffer containing 25 mM HEPES pH 7.5, 200 mM NaCl and 0.5 mM TCEP supplemented with 50U Turbo DNase and cOmplete EDTA free protease tablets, disrupted by homogenisation and sonication, and the resulting lysate clarified by centrifugation at 36,000 x *g* for 60 min at 4°C. The supernatant was applied to a 5 ml HiTrap TALON column, then washed with lysis buffer supplemented with 5 mM imidazole, before retained protein was eluted by application of the lysis buffer supplemented with 250 mM imidazole. This was applied to a 1 ml HiTrap StrepXT column, washed with lysis buffer, before elution of retained material with lysis buffer supplemented with 50 mM Biotin. The eluted material was concentrated using a Vivaspin with a 50 kDa MWCO and applied to a Superdex200increase 10/300 size exclusion column equilibrated in 10 mM HEPES pH 7.5, 150 mM NaCl, 5 % (v/v) glycerol, 0.5 mM TCEP, 0.002% (v/v) Tween-20.

**Peptide pull-down experiments**

Combinations of biotinylated SLX4 peptide corresponding to amino acids 1247-1267 (with or without Thr-1260 phosphorylated), active CDK1-CyclinB, and ATP MgCl2, were mixed and incubated at 30 deg C for 4 hours in reaction buffer 10 mM HEPES pH 7.5, 150 mM NaCl, 0.5 mM TCEP, 5 % (v/v) Glycerol, 0.002 % (v/v) Tween-20. These reactions were then mixed with StreptactinXT magnetic beads equilibrated in reaction buffer and incubated for 15 minutes at 25 deg C to allow biotinylated peptides to bind. Beads were washed with reaction buffer and subsequently GST-TOPBP1(1-290) protein was added and incubated for a further 15 minutes at 25 deg C prior to again washing with reaction buffer to remove unbound material. Samples were eluted but the addition of reaction buffer supplemented with 25 mM biotin and analysed by PAGE and Coomasie staining.

Biotin-SLX4_T1260 ‘Biotin’-GYGSEASTTDTSWLVPATPLASRSR
Biotin-SLX4_pT1260 ‘Biotin’-GYGSEASTTDTSWLVPA(pT)PLASRSR

***In vitro* fluorescence polarization**

Fluorescein-labelled peptides corresponding to amino acids 1253-1267 or 1470-1483 of SLX4 incorporating pT1260 or pT1476 respectively, were incubated at a concentration of 100 nM at room temperature with increasing concentrations of TOPBP1 WT or mutant BRCT module variants in 25 mM HEPES pH 7.5, 150 mM NaCl, 1 mM EDTA, 0.25 mM TCEP, 0.002% (v/v) Tween 20 in a black 96-well polypropylene plate. Additional experiments performed with dephosphorylated peptide were achieved though incubation with Lambda phosphatase supplemented with 1 mM MnCl2 to remove the phosphorylation on the peptide. Fluorescence polarisation was measured in a CLARIOstar multimode microplate reader. Binding curves were calculated assuming a single binding site with non-specifc binding component, using Graphpad Prism10.2.3 (GraphPad Prism, RRID:SCR_002798). Binding affinities are presented as calculated values alongside the 95% CI values. Presented graphs represent data plotted with the non-specific binding component subtracted. Each curve produced from the mean of three independent experiments, with displayed error bars representing SEM.

Flu-SLX4_pT1260 'Flu’-GYGTSWLVPA(pT)PLASRSR
Flu-SLX4_pT1476 'Flu’-GYGSPGLLDT(pT)PIRGSCT

**ALT-EJ and BIR reporter assays**

To facilitate transient depletion of CIP2A, SLX4, POLD3 or Polθ U2OS EJ2 (*1*) or U2OS BIR (*2*) cells were transfected with 30 pmol of oligonucleotides using Lipofectamine RNAimax transfection reagent (Invitrogen, 13778100), according to the manufacturer's forward transfection protocol.
crRNA (EJ2 – CTAATTACCCTGTTATCCCT, BIR - AAGATTACCCTGTTATCCCT ) targeting I-SceI recognition sequence was annealed with tracrRNA- Atto 550 (IDT) according to manufacturer’s protocol.

Cas9 (IDT, 7 pmol) and annealed crRNA–tracrRNA (8.4pmol) was incubated in 100 µl Opti-MEM for 10 min at room temperature followed by addition of 4 µl of Lipofectamine RNAimax and further incubation for 10 minutes. 250 000 cells 72 hours post siRNA transfection were then transfected with RNP complex and analysed by FACS 24 hours post transfection. The number of GFP positive cells per 30,000 (EJ2) or 10,000 (pBIR) Atto positive cells was determined and repair efficiency normalized to siCTRL samples.

**Statistical analysis**

Statistical analyses were carried out using GraphPad Prism 10 (GraphPad Software Inc.). Unpaired Student's t-test, One-way ANOVA, Two-way ANOVA, or Mann–Whitney test were used to determine statistical significance as indicated in the figure legends.

**Data availability statement**

The data generated in this study are available within the article and its supplementary data files. Data and reagents are available upon request.

**References associated with methodology.**
